# Hepatoblast iterative apicobasal polarization is regulated by extracellular matrix remodeling

**DOI:** 10.1101/2024.01.30.578046

**Authors:** Julien Delpierre, José Ignacio Valenzuela, Matthew Bovyn, Nuno Pimpao Martins, Lenka Belicova, Urska Repnik, Maarten Bebelman, Sarah Seifert, Pierre A Haas, Yannis L Kalaidzidis, Marino Zerial

## Abstract

Hepatocytes have a unique multiaxial polarity with several apical and basal surfaces. The prevailing model for the emergence of this multipolarity and the coordination of lumen formation between adjacent hepatocytes is based on asymmetric cell division. Here, investigating polarity generation in liver cell progenitors, the hepatoblasts, during liver development *in vivo* and *in vitro*, we found that this model cannot explain the observed dynamics of apical lumen formation in the embryonic liver. Instead, we identified a new mechanism of multi-axial polarization: We found that polarization can be initiated in a cell-autonomous manner by re-positioning apical recycling endosomes (AREs) to the cell cortex via fibronectin sensing through Integrin αV. Using live cell imaging we showed that this process repeats, leading to multiaxial polarity independently of cell division. We found that establishment of oriented trafficking leads to secretion of the metalloprotease MMP13, allowing neighboring hepatoblasts to synchronize their polarization by sensing extracellular matrix (ECM) distribution and enabling lumen opening. Finally, active remodeling of ECM in proximity of nascent apical surfaces closes a positive feedback loop of polarization, whereas disruption of this loop by either blocking MMP13 or downregulating Integrin αV prevents formation of the bile canaliculi network. Integration of this feedback loop into a simple mathematical model reproduces the observed dynamics of bile canaliculi network formation during liver development quantitatively. Our combined findings thus suggest a new mechanism of polarization coupling to self-organization at the tissue scale.

## Introduction

In epithelia, cells establish an apicobasal polarity, whereby the apical surfaces line a lumen, and the basal surfaces are in contact with the extracellular matrix (ECM) mediate the attachment to the underlying tissue. The cellular and molecular bases of cell polarity have been extensively studied *in vivo* and *in vitro* (Datta, 2011^1^; Blasky, 2015^2^). The apical surface of an epithelial cell is a specialized part of its plasma membrane. It has a specific protein and lipid composition and is circumscribed by tight junctions which seal the lumen (Roignot, 2013^3^; Rodriguez-Boulan, 2014^4^). Apical components are delivered through a polarized trafficking pathway and recycled by specific Apical Recycling Endosomes (AREs) (Bomsel, 1989^5^; Sheff, 1999^6^; Golachowska, 2010^7^). AREs are retained in the apical region by dynein motors moving along microtubules organized by the apically located microtubule organizing center (MTOC) (Wojtal, 2007^8^; Le Droguen, 2015^9^). In the liver, hepatocytes have a unique kind of cell polarity established during morphogenesis by the hepatic progenitors, the hepatoblasts. In contrast to cells forming columnar epithelia, hepatocytes do not have a single apicobasal axis but have a multipolar (multiaxial) organization (Treyer, 2013^10^; Gissen, 2015^11^; Morales-Navarrete, 2019^12^). This multipolarity is essential for the three-dimensional arrangement of the bile canalicular and sinusoidal networks (Morales-Navarrete, 2019^12^). The surfaces of hepatocytes contain multiple elongated apical patches which form narrow belt-like stripes. Apical patches on juxtaposed hepatocytes align to form the bile canaliculi network. Hepatocyte surfaces can also have multiple basal patches, each of which is elongated and faces one of the blood vessels of the sinusoidal network.

Key outstanding questions are: which mechanisms underlie the multi-polarity of hepatoblasts and hepatocytes? How are cell-cell interactions coordinated to generate the two complementary bile canalicular and sinusoidal networks? Early during development, at E12.5 in mice, polarization sites between juxtaposed hepatoblasts start to appear (Wood, 1965^13^). An oriented trafficking pathway is established, and the apical surfaces are then sealed by tight junctions, but the lumen is not fully opened yet, similar to the pre-apical patch (PAP) characterized in Madin-Darby canine kidney (MDCK) cells (Ferrari, 2008^14^). Later, the lumina open and expand due to secretion of osmotically active molecules by the hepatoblasts (Dasgupta, 2018^15^). The peculiarity of bile canalicular expansion is its extreme anisotropy, which is enforced by mechanical elements named apical bulkheads (Belicova, 2021^16^; Bebelman 2023^17^). This expansion results in the formation of narrow tubes. This mechanism leads to the establishment of a single polarity axis. The transition to multiaxial polarity characteristic of hepatocytes *in vivo* is not understood. Moreover, coordination at the tissue scale is required for the multiple tubular segments to be connected and form a single three-dimensional bile canalicular network. What is more, the bile canaliculi network must be interlaced with but not intersect the sinusoidal network. However, the propagation of coordination from local interactions between neighboring cells to self-organization at the tissue level is not yet understood either. Finally, the connection of the bile canaliculi network to the collecting ducts occurs one day before birth (E17), as evidenced by a strong efflux of bile reaching the gut at this time (Tanimizu, 2016^18^). However, the extent to which the bile canaliculi network is formed and dynamics of its formation before it connects to the bile ducts is also not yet characterized.

The prevailing model of bile canaliculi formation postulates that hepatocyte multiaxial polarity result from asymmetric cell divisions. Later, a switch to while symmetric cell divisions would then drive lumen expansion (Tanimizu, 2017^19^; Müsch and Arias 2020^20^). This model closely resembles the polarization model for MDCK cells and multiple other systems (Li, 2014^21^; Pohl, 2017^22^), in which cytokinesis is used to position the nascent apical surface at the middle of the cell-cell interface. Then the midbody guides the orientation of the trafficking through ARE positioning (Wakabayashi, 2005^23^; Mangan, 2016^24^) and leads to the establishment of tight junctions (Kojima, 2001^25^; Wang, 2014^26^). In contrast to MDCK cells which divide symmetrically in such a way that both daughter cells share the previously established lumen, hepatoblasts could divide asymmetrically, resulting in the existing lumen being inherited by one daughter cell and a second lumen being formed at the cytokinesis site. As a result, one cell becomes multipolar (Rodriguez-Fraticelli, 2010^27^; Lázaro-Diéguez, 2013^28^; Tanimizu, 2017^19^). It has been proposed that hepatoblasts divide asymmetrically up to E17.5 and then switch to symmetric divisions to connect the lumina into a network (Tanimizu, 2017^19^). This model could thus explain the multipolar organization of hepatocytes and the coordination of lumen formation between juxtaposed hepatocytes. However, the basic assumptions of the model have never been tested *in vivo*. Moreover, it has been shown that adult primary hepatocytes can form multiaxial polarity and thin tubular lumina *in vitro* (Mooney, 1992^29^; Zeigerer, 2017^30^; Kaur, 2023^31^), which calls for a refinement of this cell division-centered model. Additionally, the key element for hepatocyte polarization *in vitro* is their contact to ECM (Dunn, 1989^32^), which is required for polarization in multiple systems (Yu, 2005^33^; Bryant, 2014^34^) and sufficient to orient intracellular trafficking (Ojakian, 1997^35^; Akhtar, 2013^36^; Li, 2016^37^). Hepatoblasts express high levels of ECM components such as fibronectin and vitronectin (Sugiyama, 2013^38^). However, how ECM sensing contributes to the synchronous and precise positioning of apical surfaces between juxtaposed polarizing hepatocytes is unclear.

To address these outstanding questions, we investigated the mechanisms of generation of hepatocyte multipolarity both by quantitative analysis of tissue during liver development *in vivo* and using primary hepatoblasts differentiating and polarizing *in vitro*. Our study reveals a new mechanism of multiaxial polarization of hepatoblasts, which ensures synchronized polarization of juxtaposed hepatoblasts and tissue-level self-organization through dynamic ECM sensing and remodeling.

## Results

### Lumen generation during development cannot be explained by hepatoblast divisions alone

To understand the mechanisms underlying hepatocyte multiaxial polarity *in vivo*, we performed an accurate quantification of apical lumen formation in the developing mouse liver. As a first step, we set to establish a detailed timeline of events leading to bile canaliculi network development, by quantifying the formation of lumina. We collected embryonic livers at multiple developmental stages and performed high-resolution deep tissue imaging of 100 μm optically cleared sections immunostained with a set of subcellular markers. We developed an image analysis pipeline for unbiased quantification of apical lumina. Hepatocyte apical lumina are actin-rich structures. We confirmed by electron microscopy (EM) that they are as small as 1 μm^3^, lined by tight junctions, and display the aminopeptidase CD13 on their surface (Feracci, 1987^39^) (Figure 1A). Therefore, we segmented apical lumina defined as discrete structures of at least 1 μm^3^, positive for CD13, the cell cortex marker (F-Actin), and the tight junction marker Zonula Occludens 1 (ZO1) (Figure 1B, C).

**Figure1:**
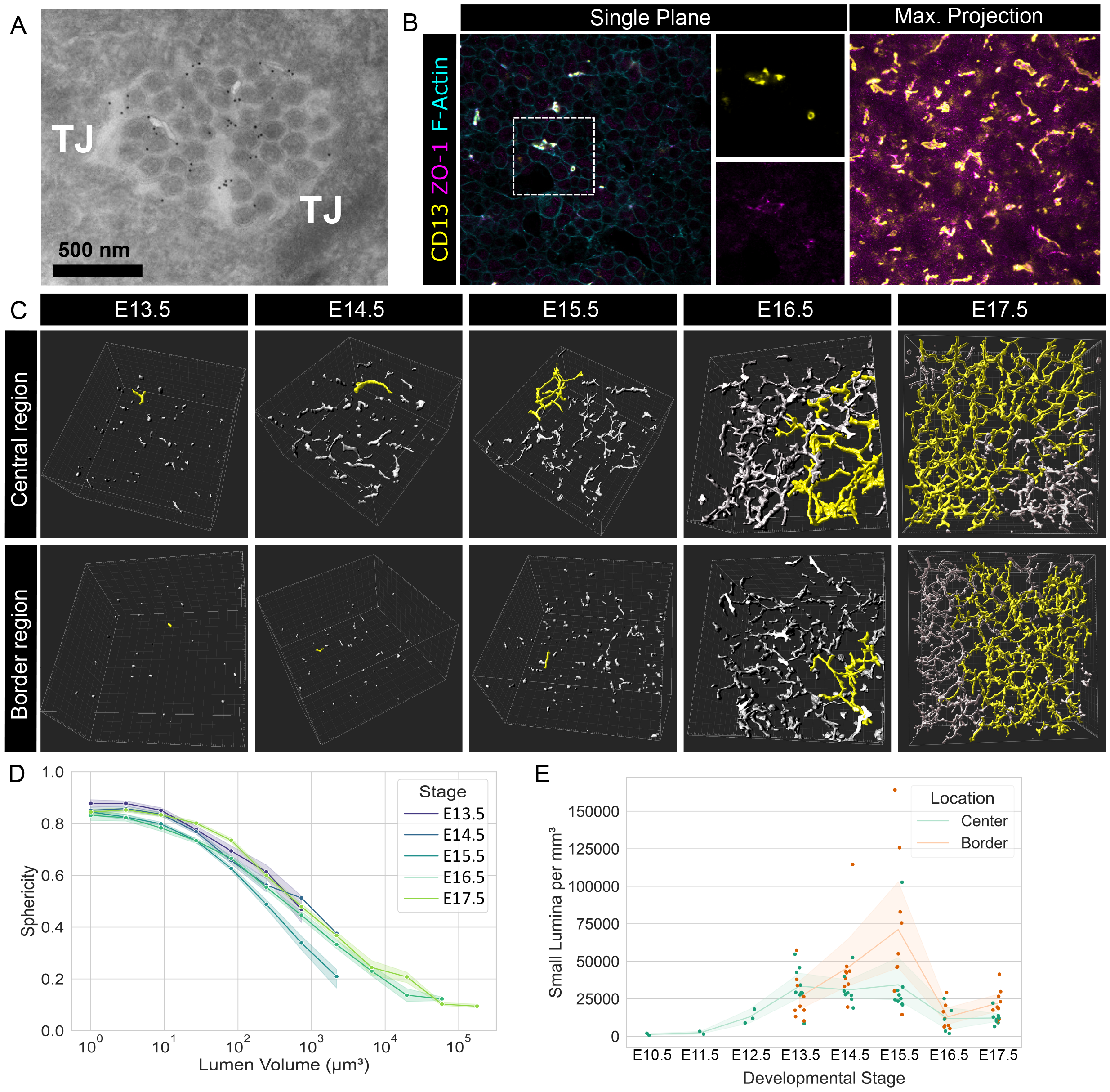
Timeline of bile canaliculi network formation. A: Electron micrograph of a liver section at E14 stained with Immunogold directed against CD13. Lumina is opened and filled with microvilli, CD13 is visible in the luminal space. TJ: Tight junctions. B: Single plane image and maximum projection over 50 μm of immunostaining directed against luminal markers (CD13, ZO-1) and a cell border marker (F-actin labelled with Phalloidin), on fixed liver section at E14.5 in the hilum region. Dashed white square shows the region selected for single channel illustrations. Scale bar: 10μm C: 3D reconstruction of apical surfaces (CD13+, ZO-1+, F-actin+) of liver sections, in the central region and in the periphery of the organ, between the initial lumen formation and network formation. The white box shows the image frame, ticks are 10μm. The largest continuous object is shown in yellow. The field of view at E17 is larger (in X and Y) than the other timepoints. D: Quantification of the relation between lumina sphericity and volume. Data has been pooled to the lower bin and each bin is represented by a point. Minimum bin size is 5. Sample size: E13.5=994, E14.5=1719, E15.5=2619, E16.5=1578, E17.5=2884. E: Quantification of the amount of small spherical lumina (<10μm) density across development. Green curve shows the central region (close to the hilum) and orange curve shows the border region (at the periphery of the organ). The graph shows the same dataset as D. Samples where no lumina have been found are not shown (one at E10.5 and one at E11.5). Sample size per region: E10.5-E12.5= 3, 13.5-17.5=9. For all plots, shaded area corresponds to 95% confidence interval. D and E are extracted from the same dataset of images.

First, we found strong differences in the number of lumina across frontal sections of the organ (Figure S1A, B). Specifically, the bile canaliculi network at the central region close to the hilum appeared to be further developed than the peripheral regions (Figure 1C). This is similar to findings of inhomogeneity of bile duct development (Takashima, 2015^40^; Tanimizu, 2016^18^). We then found that, initially, lumina appeared as small spherical objects, which later turned into tubes. Such a transition can be quantified as a decrease in sphericity. We found that lumen sphericity drops only after its volume reached 10 μm^3^ (Figure 1D). Since small spherical lumina were detected at all time points analyzed and in all considered regions (Figure 1E), we deduce that hepatoblasts continuously generate new lumina across the tissue. Hepatoblast divisions also occur continuously throughout the tissue, but it is not clear whether they are able to account fully for the number of lumina observed. Therefore, we decided to investigate the evolution of lumen number quantitatively.

The number of lumina is the result of two main processes: formation of new lumina and lumen fusion. At later stages of development, lumen fusion dominates, progressively resulting in all lumina connected into a single bile canalicular network in adult liver (Morales-Navarrete, 2019^12^). The number of distinct lumina increased from E12.5, peaked at E15.5, and declined between E15.5 and E17.5 (Figure 2A). Such a decrease of lumen number could be due to resorption of lumina or their fusion. Since the percentage of tissue occupied by apical surfaces kept increasing without any sign of saturation between E15.5 and E17.5 (Figure 2B), we conclude that fusion must be the main reason for the decreasing number of lumina. Notably, we could identify clearly multipolar hepatoblasts at E14 (Figure 2C) and some instances of tight junctions connecting distinct lumina (Figure 2D). In summary, generation of new lumina by hepatoblasts dominates between E12.5 and E15.5, whereas fusion between lumina takes over after E15.5. Therefore, our results suggest that predominant lumen fusion and network formation begins at E15.5, two days earlier than previously estimated (Tanimizu 2016^18^). Interestingly, although the initiation of lumen formation in the central and peripheral zones are very different, the transition from increase to decrease in the number of lumina occurs at the same time (E15.5) (Figure 2A, B), suggesting that the fusion process is coordinated at the tissue scale.

**Figure2:**
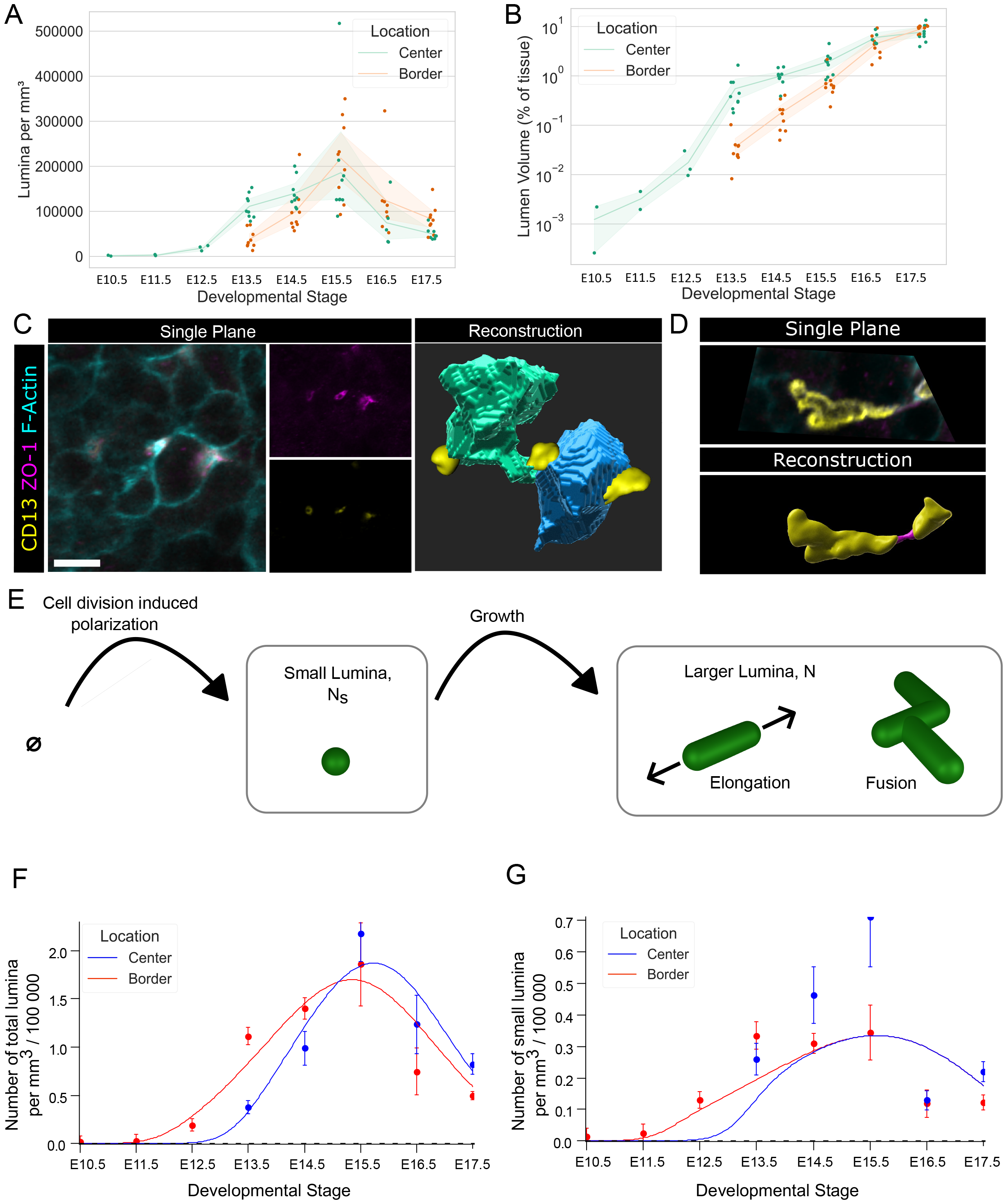
Lumen generation exceeds cell division. A-B: Quantifications of lumina parameters across development. Green curve shows the central region (close to the hilum) and orange curve shows the border region (at the periphery of the organ). C: Illustration of multipolar hepatoblasts. Immunofluorescence image of lumina markers (ZO-1, CD13) and cell border (F-Actin labelled with Phalloidin). Smaller images show individual channels. Reconstruction of the two multipolar cells visible on the single plane images. Scale bar = 10μm D: Illustration of junctional bridge connecting two distinct apical surfaces. Immunofluorescence image of lumina markers (ZO-1, CD13) and cell border (F-Actin labelled with Phalloidin). Reconstruction of the two lumen in yellow and the junction bridge in magenta. E: Schematic of the *in silico* model. Cell division leads to the formation of small lumina. Growth leads to elongation and fusion. F-G: Comparison between the simple model and the quantification. Curves shows the model prediction Points and error bars corresponds to the data. Central area is shown in blue and the border area is shown in red. For all plots, shaded area corresponds to 95% confidence interval. A and B are extracted from the same dataset of images as figure 1D-E, samples where no lumina have been found are not shown (one at E10.5 and one at E11.5), sample size per region: E10.5-E12.5= 3, 13.5-17.5=9.

In light of these findings, we investigated whether cell division alone could explain the observed lumen formation across the organ. To do so, we used published data on hepatoblast proliferation during liver development (Weiss, 2016^41^) and our previous quantifications of lumina formation to feed a simple quantitative model (Appendix). Its parameters were fitted to experimental data. We distinguished two classes of lumina: 1) small spherical lumina, which most probably correspond to newly formed lumina (primary polarization); 2) large elongated and branched lumina (Figure 2E). The latter population grows by maturation of the primary spherical lumina. These lumina also elongate and fuse. Since, we cannot perform direct *intravital* measurements of the rates of lumen formation and fusion, we inferred them from experimental data by the following procedure: first, we described the dynamics of lumen number and volume by a set of ordinary differential equations (ODEs) (Appendix). In order to formulate these ODEs, we assumed that (a) each hepatocyte division could lead to formation of a new lumen; (b) lumen elongation and fusion have similar behaviors regardless of the region of the tissue; (c) the elongation and fusion rates are proportional to the number of “free ends” of lumina (Appendix). The parameters in these ODEs were either directly taken from the experimental data (e.g., the number of new lumina generated after the onset of lumenogenesis was set to the number of hepatoblast divisions during liver development from E12.5 to E17.5 as previously reported in Weiss, 2016^41^), or were globally fitted to the experimentally observed dynamics of lumen number and volume (e.g. elongation and fusion rates were fitted in this way). The parameters best fitting the data correspond to each cell division event leading to the formation of an additional lumen. Even though this yields the maximum number of lumina that could be generated by this mechanism, the model could only explain 75% of the experimentally observed small lumina (Appendix) and failed to recapitulate the experimentally observed peculiar dynamics of lumina density even qualitatively (Figure 2F, G). From these results, we conclude that hepatoblast polarization must also rely on a mechanism independent of cell division.

### Multiple AREs can be observed within the same cell prior to multipolarization

To elucidate cell-division-independent mechanisms of the establishment of multiaxial polarity, we explored the establishment of polarized trafficking by following the intracellular localization of AREs at different stages of liver development. AREs are a cluster of vesiculo-tubular compartments (Goldenring, 2015^42^) characterized by the presence of recycling endosomal markers such as Rab11 and apical surface proteins. Since the tubules of AREs cannot be resolved by conventional light microscopy, we visualized and segmented AREs as single objects co-stained with Rab11 and CD13 antibodies in thick tissue samples (Figure 3A). Depending on their localization, AREs appeared either as perinuclear spherical objects or flat patches next to the plasma membrane. This morphological change can be quantified as a reduction in ARE sphericity (Materials & Methods). At E12.5, CD13 was located both at the cell surface of hepatoblasts and intracellularly within the AREs. This polarization stage corresponds to the inverted polarity described for MDCK cells (Ferrari, 2008^14^). One day later, at E13.5, most CD13 relocated to the AREs and was hardly detectable at the cell surface. At E13.5 and E14.5, most AREs were recognizable as single flat clusters near the cell border, characterized by a low sphericity (Figure S2A and B). In contrast to MCDK cells, where the polarization includes formation of the PAP, (i.e. ZO-1 positive plasma membrane patches with AREs located nearby), in hepatoblasts the AREs were located close to the plasma membrane and mostly negative for ZO-1 (Figure S2C), suggesting that their localization does not depend on the existence of ZO-1 patches at the cell surface. Surprisingly, at E15.5, we could discern two distinct populations of AREs within single hepatoblasts, one subapical and one located away from the lumen and in proximity to a different region of the cell border. This configuration could still be observed at E16.5, even in multipolar hepatoblasts, suggesting that the process of polarization is not yet complete (Figure 3A). Therefore, the process of polarization of hepatoblasts comprises the splitting of subapical AREs and the relocation of a second ARE population to the position of a new putative apical pole (Figure 3B).

**Figure3:**
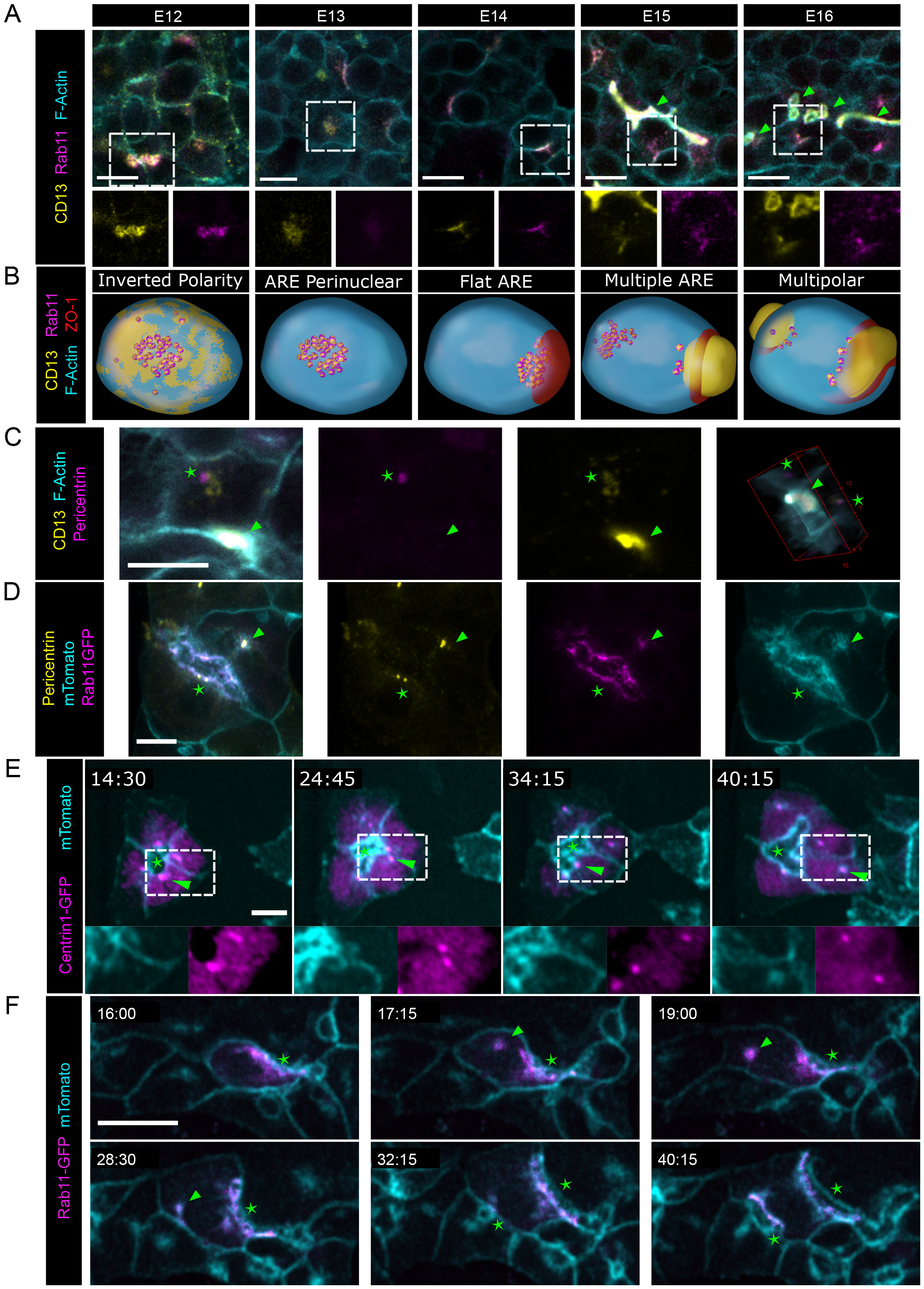
AREs and centrosomes split and are re-located away from lumina after hepatoblast polarization. A: ARE locations in vivo. Single plane images of liver sections at various stages immunostained against ARE markers (Rab11, CD13) and cell border (F-actin labelled with Phalloidin). Arrowheads point established lumina. Dashed squares show the magnified areas. Magnified areas show individual channels. B: Schematic of the steps leading to hepatoblast multipolarization observed in A. C: Centrosome location in polarized hepatoblasts. Single plane image of liver sections at E14.5 immunostained against apical (CD13), centrosomal markers (Pericentrin) and cell border (F-actin labelled with Phalloidin). 3D view of the same lumen. Arrowheads point established lumina and stars highlights centrosomes. D: Immunofluorescence image of fixed primary hepatoblasts expressing a membrane targeted tdTomato (mTomato), transfected with mRNA coding for Rab11 fused with EGFP and stained against a centrosomal marker (Pericentrin). Arrowhead shows a centrosome located away from a lumina together with pAREs. Star shows a sub-apical centrosome. E-F: Fluorescence image of sequential timepoints of live imaging in vitro. Primary hepatoblasts expressing mTomato were transfected with mRNA (coding for either Rab11 or Centrin1 fused with EGFP). Timestamp units = hh:mm. Arrowheads point the re-locating population of AREs. Stars labels the established lumina. Scale bar: 10μm

AREs are clustered around the centrosome or MTOC through a dynein-dependent mechanism (Perez Bay, 2013^43^). The presence of multiple clusters of AREs could be explained either by the duplication of the centrosome or by the presence of a MTOC without centrosome. We therefore stained embryonic liver tissue sections with antibodies directed against CD13 and the centrosome marker Pericentrin. We found that all non-dividing hepatoblasts had only one centrosome and, when the AREs split and the second population of AREs relocated away from an established lumen, they had the centrosomes in close proximity (Figure 3C). We conclude that splitting of AREs is not accompanied by centrosome duplication.

### Apical recycling endosome splitting and movement lead to formation of multiple lumina

In hepatoblasts both *in vitro* and *in vivo*, centrosomes were always surrounded by Rab11-positive AREs located either near a lumen or in a second location distant from the primary lumen (Figure 3D). The establishment of a second oriented trafficking axis in an already polarized hepatoblast would imply that both AREs and centrosome must relocate from the primary lumen to a second, new putative polarization site. To test this hypothesis, we performed live cell imaging on isolated primary hepatoblasts *in vitro* (Belicova, 2021^16^). We took advantage of transgenic mice expressing the membrane-targeted fluorescent protein tandem-dimer Tomato (mTomato) to visualize membrane dynamics on isolated hepatoblasts. We transiently transfected the hepatoblasts with mRNA encoding the centrosome marker Centrin-1 tagged with GFP (Centrin-1-GFP). The presence of mTomato on the AREs allowed us to follow this endosomal compartment (Figure 3D). We could directly visualize the movement of centrosomes labelled by Centrin-1-GFP across the hepatoblast together with a population of ARE vesicles reaching a different region of the plasma membrane (Movie 1 and Figure 3E). We refer to this moving population of AREs as peri-centrosomal AREs (pAREs) to emphasize its association with the centrosome.

To confirm that the endosomal compartment which splits into two populations are the AREs, we transfected hepatoblasts with mRNA encoding a GFP-tagged Rab11 and followed the dynamics of AREs during secondary apical pole formation. As expected, following the initial polarization, most AREs co-labelled with GFP-Rab11 and mTomato were located sub-apically (Movie 2 and Figure 3F). However, after some time, AREs split into two sub-populations, one of which moved away from the initial lumen. This second cluster reached another part of the cell where it persisted for a few hours. Eventually, a new lumen formed at that site. This confirms our hypothesis that multiaxial polarity can result from iterative splitting and re-positioning of AREs without involvement of cell divisions. This mechanism could also explain the observed multiaxial polarization of postmitotic adult hepatocytes *in vitro*.

Lumen formation and expansion require sealing of apical membranes of neighboring hepatoblasts and therefore the coordination of their polarization processes. We observed that the positioning of pAREs in neighboring hepatoblasts is not synchronous (Movies 1 and 2). Instead, one cell chooses a polarization site by positioning its pAREs, and the neighboring hepatoblast responds after a few hours. We therefore searched for potential cues guiding pAREs re-location and spatial synchronization.

### AREs reach the cell border in a cell-autonomous manner

Cell polarization occurs in two steps: first, the establishment of oriented trafficking and, second, the formation of a lumen. In MDCK cells, establishment of oriented trafficking through ARE positioning is associated with the formation of a PAP, which provides localization signals through the Cingulin-ZO-1-Occludin complexes (D’Atri, 2002^44^; Mangan, 2016^24^). However, in hepatoblasts, the localization of pAREs occurs before ZO-1 recruitment (Figure S2C), implying that the mechanism of pAREs positioning must be different in these cells. To assess the importance of tight junctions in oriented trafficking in hepatoblasts, we silenced ZO-1 by transfecting primary hepatoblasts with siRNA. In control cells, AREs could be found in proximity of Occludin-positive plasma membrane patches. Upon ZO-1 knockdown (KD), AREs were still localized in proximity of the plasma membrane, but they did not colocalize with Occludin patches (Figure 4A). As a result, overall cell polarization was impaired (Figure S3A, B). This suggests that interactions with tight junctions are not required for AREs to position themselves near the plasma membrane but are required for localizing AREs proximally to tight junctions.

**Figure4:**
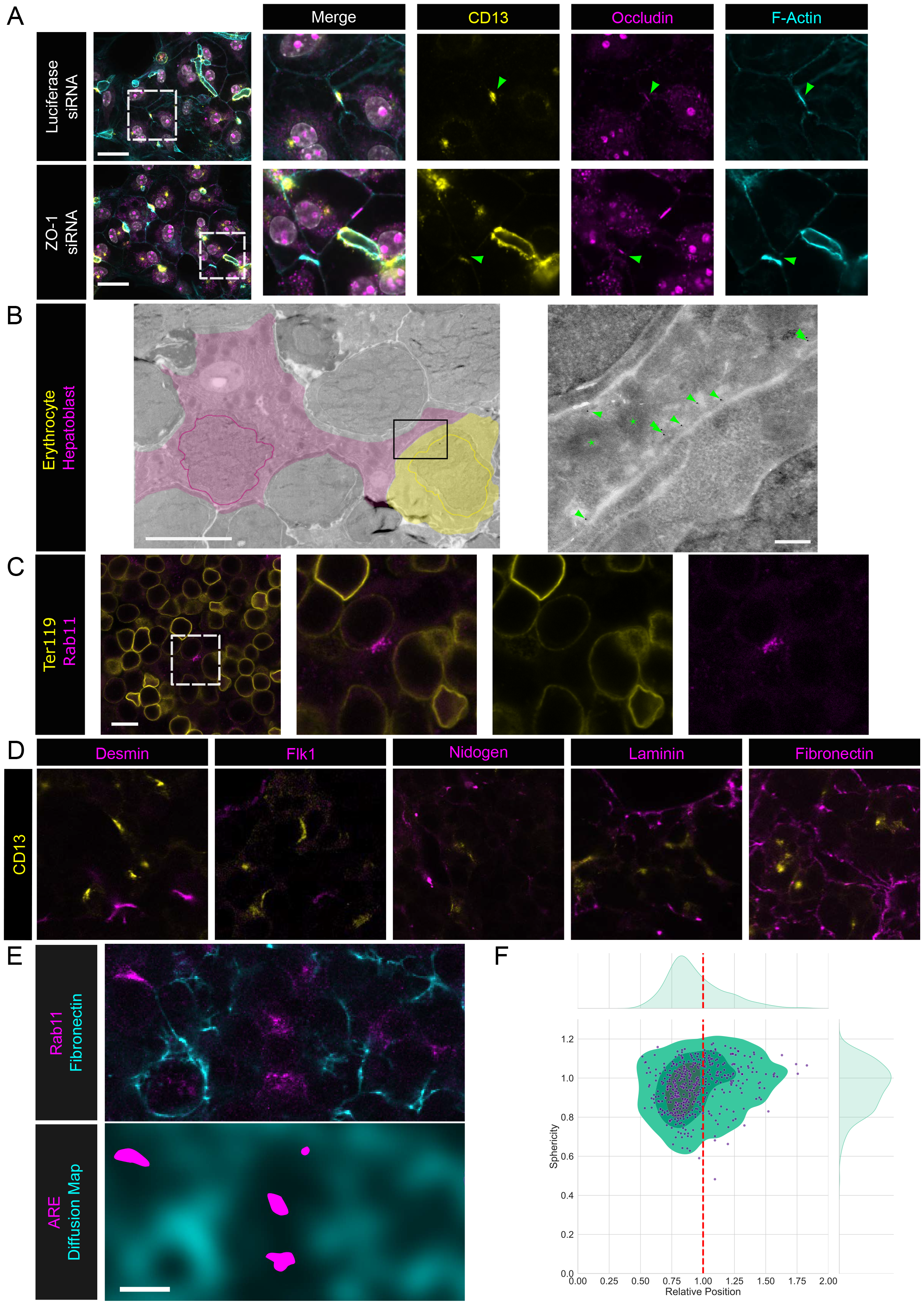
pAREs reach the cell border in a cell autonomous manner and are located away from the ECM. A: Single plane images of immunostaining directed against PAP markers (CD13, Occludin) and a cell border marker (F-actin labelled with Phalloidin), on fixed primary hepatoblasts, after treatment with siRNA directed against Luciferase (non-targeting) or ZO-1. Arrowheads point pAREs located close to the cell border in both conditions. B: Electron micrographs of E14 liver sections stained with immunogold directed against CD13. Black square shows the magnified area. Colors: Magenta=Hepatoblast Yellow=Erythropoietic progenitor. Colored lines highlight nuclei contour. Arrowheads point gold particles. Stars mark the centrioles. Overview image scale bar: 5μm; Magnified image scale bar: 500nm. C: Single plane image of E14 liver section immunostained against erythropoietic progenitor marker (Ter119) and AREs marker (Rab11). The dashed square shows the magnified area. Scale bar: 10μm D: Single plane images of a liver sections at E13.5 immunostained against CD13 and various basal markers. E: Single plane image of a liver section at E13.5 immunostained against ARE (Rab11) and ECM (Fibronectin) markers and the same region showing the segmented ARE and the diffused ECM signal. F: Quantification of the relative position against the sphericity of AREs. Points represent individual AREs. Green shaded areas represent the kde levels of the points distribution. kde is projected along each axis. Red dashed line is highlighting the relative position of 1. Scale bar is 10μm unless stated otherwise.

Our results further suggest that pARE positioning occurs asynchronously between neighboring cells. To verify this observation, we examined the sub-cellular distribution of AREs *in vivo* by EM. Thin sections were stained with immunogold directed against the apical marker CD13 to identify AREs (Figure 4B). We found pAREs located at the cell border of hepatoblasts in regions without junctions, confirming that pARE positioning near the plasma membrane can occur in the absence of junctions. Furthermore, the neighboring cell did not display any sign of oriented trafficking and, what is more, it appeared morphologically to be an erythropoietic progenitor. Indeed, erythropoietic progenitors are abundant in the embryonic liver at these stages of development (Moore and Metcalf, 1970^45^; McGrath, 2008^46^). To validate this interpretation, we performed immunostaining against Rab11 and Ter119, an erythropoietic progenitor marker (Figure 4C). Indeed, we found pAREs located at cell borders neighboring Ter119-positive erythropoietic progenitors. These results suggest that pARE positioning can occur in the absence of junctions and in a cell-autonomous manner, i.e. independently of the presence of a neighboring hepatoblast.

### AREs are located away from the ECM

In MDCK cells, ECM-derived signals determine the orientation of cell polarization through Integrin β1 (Yu, 2005^33^; Bryant, 2014^34^). AREs localize near the apical plasma membrane, providing a hallmark of the formation of an apical pole (Bryant, 2014^34^). In isolated hepatocytes, ECM provides basal signals, which in combination with cell-cell junctions (E-Cadherin) are sufficient for the sub-apical positioning of AREs and to induce cell polarity (Bryant, 2014^34^, Zhang, 2020^47^). Therefore, to characterize cell polarization in the developing liver tissue, we explored the distribution of several ECM proteins and other potential basal signal sources, as well as AREs. First, we performed immunostaining against CD13 and multiple basal signal candidates and qualitatively evaluated their distribution (Figure 4D, S4). Whereas Laminin, Collagen IV, and blood vessels were sparsely distributed and many hepatoblasts were not in contact with them, Fibronectin had a broader distribution across the liver parenchyma as described previously (Shiojiri, 2004^48^). Therefore, Fibronectin is a promising candidate ECM protein that could provide a signal for hepatoblasts to polarize.

Hence, we sought to correlate the spatial distribution of Fibronectin with the location of AREs, as they localize early to the presumptive apical region in the polarization process. Therefore, we decided to test the correlation of ARE positioning and Fibronectin density in the vicinity of AREs. Accordingly, we imaged volumes of liver tissue stained against Fibronectin and Rab11 (as a marker for AREs, Figure 4D). Since AREs are intracellular, whereas Fibronectin is extracellular, simple signal overlap cannot be used to quantify their co-localization. In order to correlate the distance of the Fibronectin signal with ARE position, we created a computational proxy for inverse distance from the Fibronectin localization (Materials and Methods): we performed *in silico* simulation of anisotropic diffusion of the Fibronectin signal into the cell interior. This anisotropy strongly favored diffusion along the cell borders to preserve the local inhomogeneity of Fibronectin signal density and overall cell morphologies (Figure 4E). We measured the mean intensity of the diffused signal within the segmented AREs (as compact clusters of endosomes) (n=465) and normalized it by the intensity in the surrounding area. We called this normalized intensity “relative position” because the values below 1 and above 1 correspond to localization of AREs away from or near Fibronectin, respectively. Most relative position values were found to be below 1, which suggests that, overall, AREs are positioned farthest away from fibronectin. As AREs close to the plasma membrane are flatter, the sphericity of AREs could be used as a numerical proxy for their distance from the cell surface. We found that a low sphericity was associated with a lower value of relative position (Figure 4F). Altogether, these results suggest that AREs are positioned to the cell border away from Fibronectin, and thus support the hypothesis that, as in MDCK cells, ECM signaling is involved in establishing polarized trafficking in hepatoblasts *in vivo*.

### MMP13 activity and Integrin αV signaling are required for hepatoblast polarization

Since hepatoblasts must set up multiple apical poles and because apical pole position could be determined by ECM distribution, we tested whether hepatoblasts could conversely remodel Fibronectin concomitantly with multiaxial polarization. For this, we co-stained thick sections of embryonic livers with antibodies directed against Fibronectin and CD13, and quantified Fibronectin intensity distribution relative to the apical lumen of hepatoblasts at the onset of polarization at E13.5 and at the onset of network formation at E16.5. Based on this staining, we drew a distance map from CD13, encompassing both apical surfaces and AREs (Figure 5A), and quantified the Fibronectin intensity as a function of the distance from CD13 at each time point. We found that the fibronectin intensity distribution shifted away from CD13 over time (Figure 5B). Such a shift can be explained either by increased ECM deposition on the basal side or by removal from the apical side. The latter mechanism seems more plausible, given that at E13.5 Fibronectin was almost homogeneously distributed across the parenchyma.

**Figure5:**
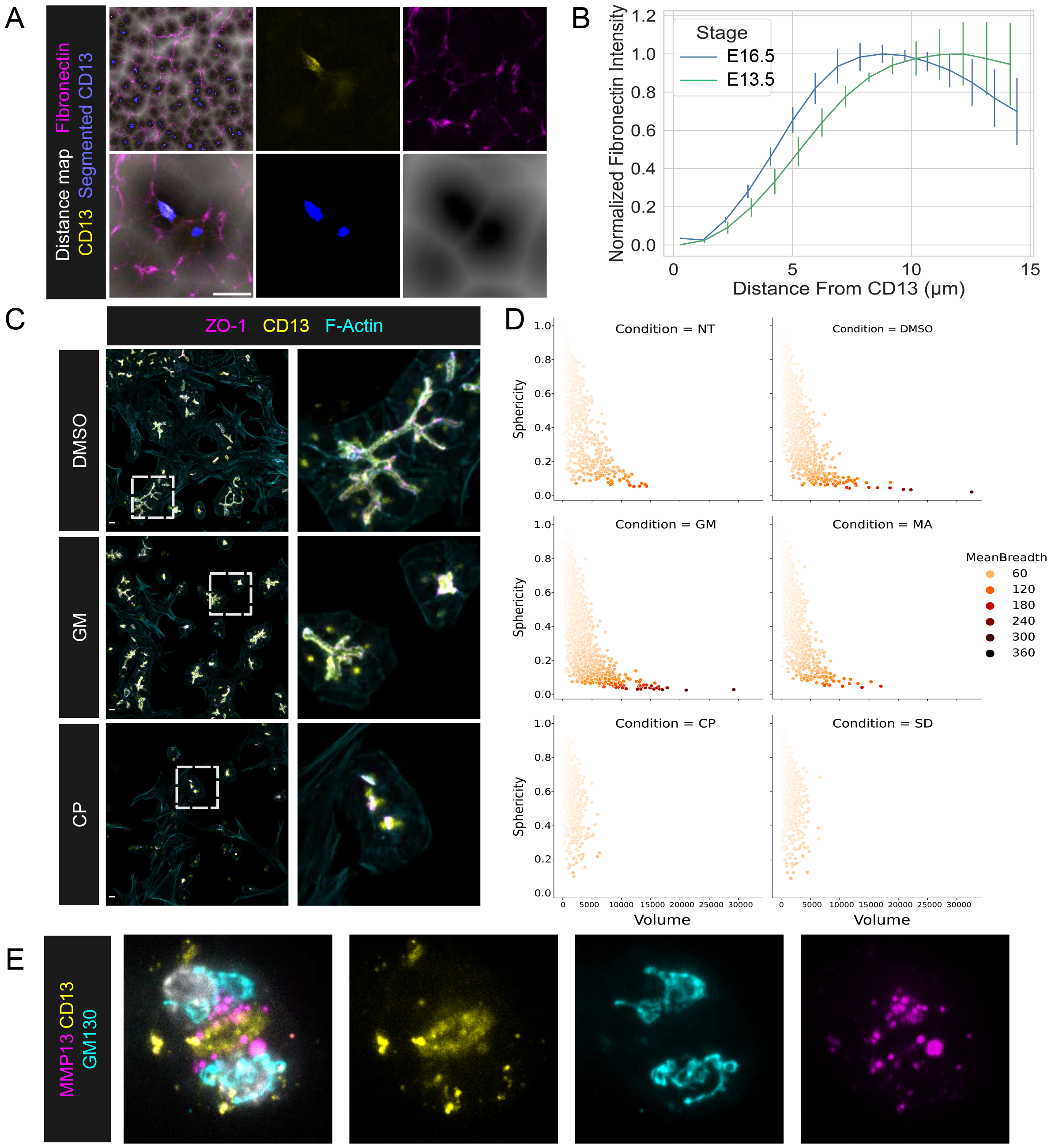
MMP13 is required for hepatoblast polarization. A: Single plane image of E13 liver section immunostained against apical (CD13) and ECM (Fibronectin) markers overlayed with the processed image: segmented CD13 and the diffusion map build with the segmentation results. B: Quantification of fibronectin intensity as a function of the distance from CD13 at the onset of lumen formation (E13) and network formation (E16). Normalized mean values of 3 images per timepoint. C: Single plane images of fixed primary hepatoblasts treated with solvent (DMSO) or mmp inhibitors (GM and CP) and immunostained against lumina (ZO-1, CD13) and cell border (F-actin labelled with Phalloidin) markers. D: Quantification of lumina morphology (sphericity as a function of volume and mean breadth color coded) changes upon treatment with mmp inhibitors. Points correspond to lumina. E: Single plane image of fixed primary hepatoblasts immunostained against MMP13, a golgi marker (GM130) and apical marker (CD13). The star labels the lumina. GM: GM6001; CP: CP471474; MA: Marimastat; SD: SD 2590 hydrochloride; Scale bars: 10μm

To test this hypothesis, we investigated the contribution of ECM degradation by metalloproteases to the polarization process in isolated primary hepatoblasts. We listed the metalloproteases expressed during polarization at E14.5 based on the GXD database (Smith, 2019^49^), and selected 4 metalloprotease inhibitors with various substrate specificities and complementary targets among the expressed ones (Figure S5). We found that 2 out of 4 inhibitors blocked lumen formation at the PAP stage (Figure 5C, D, and S5). Their target specificity and pattern of expression (Figure S5) suggest matrix metalloprotease 13 (MMP13) activity is required for lumen formation. To validate this interpretation, we performed immunostaining against MMP13 and found that the intercellular pool of MMP13 is located in large sub-apical vesicles (Figure 5E), indicating polarized secretion of MMP13. Notably, it was reported previously that MMP13 effectively cleaves Fibronectin (Zhang, 2012^50^). Along with the correlation of ARE location away from fibronectin, this result therefore suggests that blocking the removal of Fibronectin, i.e. by MMP13 inhibition, prevents apical surface formation. These results further suggest that similar to other systems (Iruela-Arispe, 2009^51^; Akhtar, 2013^36^), Fibronectin acts as a negative cue in the process of establishing multiaxial polarization also in hepatoblasts.

If Fibronectin acts as a negative cue, Integrin sensing of Fibronectin should play a role during multiaxial polarization. To test this, we knocked down Integrin αV, known to be involved in Fibronectin signaling (Morgan, 2009^52^). The KD of Integrin αV affected the morphology of cells by increasing cell spreading and preventing proper polarization (Figure 6A, B). Consequently, hepatoblasts did not form normal lumina but instead localized ZO1 and CD13 in Phalloidin-positive small spherical cytoplasmic structures. These structures could well correspond to the previously reported vacuolar apical compartment (VAC) that forms when the delivery of apical surface proteins is delayed in MDCK cells (Vega-Salas, 1987^53^; Martin-Belmonte, 2008^54^). They were more frequent than lumina in the controls (Figure 6B), and in multiple cases were located intracellularly near the ECM (Figure 6C). Therefore, we conclude that Integrin αV is required for establishment of proper polarized trafficking.

**Figure6:**
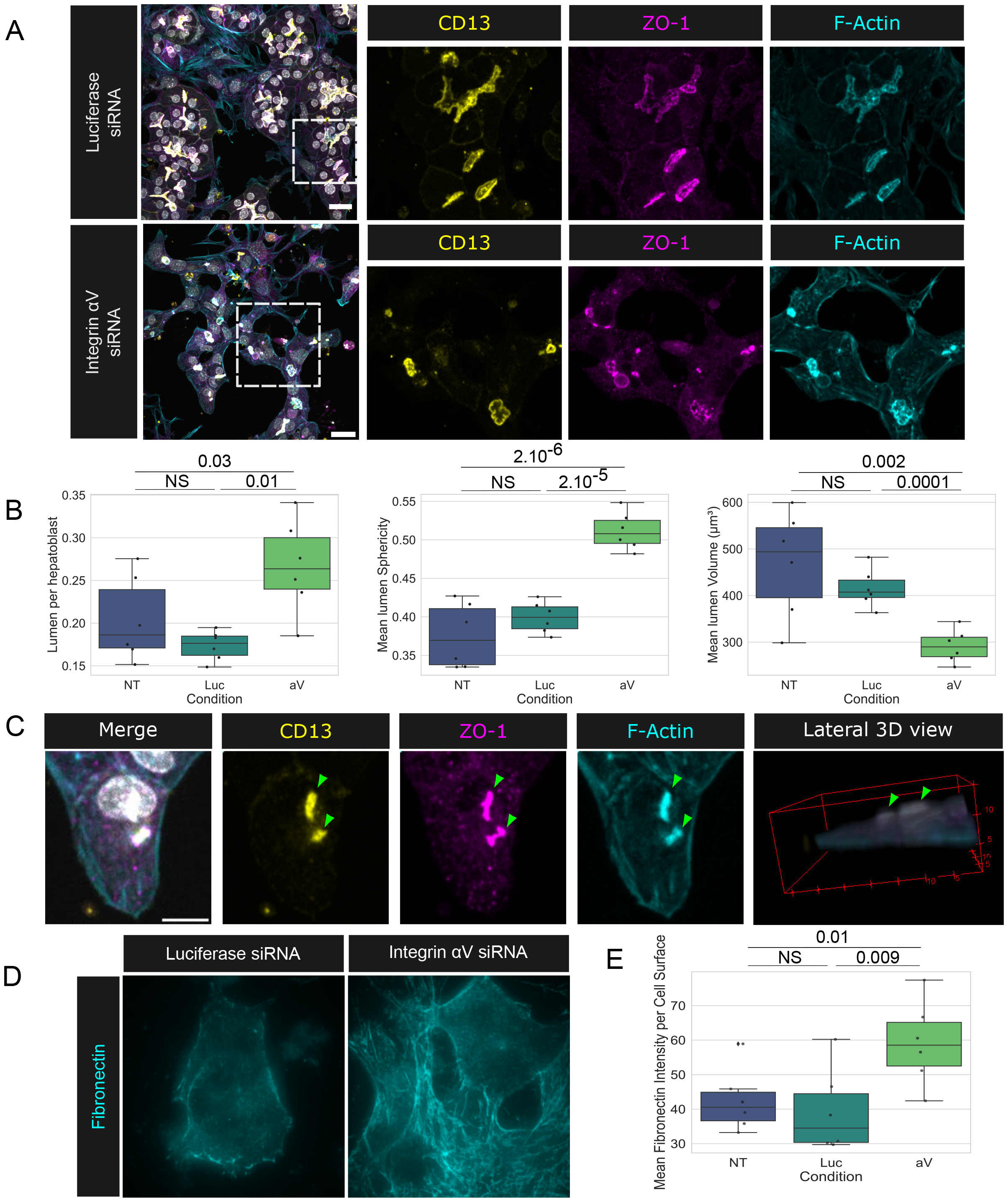
Integrin αV is required for hepatoblast polarization. A: Single plane images of fixed primary hepatoblasts treated with either luciferase (non-targeting) or Integrin αV siRNA and immunostained against lumina (ZO-1, CD13) and cell border (F-actin labelled with Phalloidin) markers. Dashed boxes show the magnified area. B: Quantifications of Integrin αV KD effect on polarization. Points are showing mean values from 20 images per well. P-values are indicated when lower than 0.05. C: Single plane image and 3D view of fixed primary hepatoblasts treated with Integrin aV siRNA and immunostained against VAC (Intracellular, ZO-1, CD13) and cell border (F-actin labelled with Phalloidin) markers. Arrowheads points VACs. D: Single plane image of fixed primary hepatoblasts treated with Luciferase (non-targeting) or Integrin αV siRNA and immunostained against Fibronectin. E: Quantification of fibronectin intensity changes upon Integrin αV KD. Points represent mean values of 20 images. P-values are indicated when lower than 0.05. Scale bars: 10μm

Given that MMP13 activity is required for hepatoblast polarization, one might expect that KD of Integrin αV impacts ECM remodeling. Although in our culture system hepatoblasts were seeded in diluted Matrigel containing only trace amounts of Fibronectin not detectable by immunostaining, we detected strong Fibronectin signal around hepatoblasts (Figure 6D). This observation is in agreement with previous findings showing that hepatoblasts secrete Fibronectin *in vitro* (Tamkun, 1983^55^; Odenthal, 1992^56^; Chagraoui, 2003^57^). As expected, KD of Integrin αV led to an increase of Fibronectin signal (Figure 6E). This suggests a relationship between ECM sensing and remodeling.

### A positive feedback loop of polarized trafficking, ECM degradation and lumen formation explains the observed lumen dynamics

Our results led us propose the following model of hepatoblast polarization: 1) ECM guides oriented trafficking through Integrin αV prior to polarity establishment; 2) once oriented trafficking is established, hepatoblasts further degrade the ECM locally at the presumptive apical site, thus providing a cue for a potential polarization site to the neighboring hepatoblasts; 3) neighboring hepatoblasts sense the lack of ECM and respond by orienting polarized trafficking towards the same site. This whole process constitutes a positive feedback loop that facilitates the formation of secondary lumina after polarity is established. Such a feedback loop can be expected to impact the dynamics of bile canaliculi network formation. Since the lumen formation model based on cell division did not explain our experimental data, we added to this model the possibility of spontaneous polarization and induced secondary polarization (i.e. formation of a second polarity axis) through this feedback loop (Figure 7A, Appendix). This model was able to explain the observed number of lumina (Figure 7B), formation of small lumina (Figure 7C), and lumen volume (Figure 7D). It allowed us to infer the lumen generation rate (Figure 7E). We found that including the positive feedback loop in the model significantly improves the quality of the fit of the model to experimental data, even considering the increased number of parameters (Appendix). Interestingly, the model predicts that the positive feedback loop leads to formation of a secondary wave of polarization (Figure 7E and Appendix), which can be interpreted as acquisition of multiaxial polarity. In summary, our model supports the hypothesis that a positive feedback loop involving ECM deposition and remodeling by hepatoblasts controls their multiaxial polarization (Figure 7F).

**Figure7:**
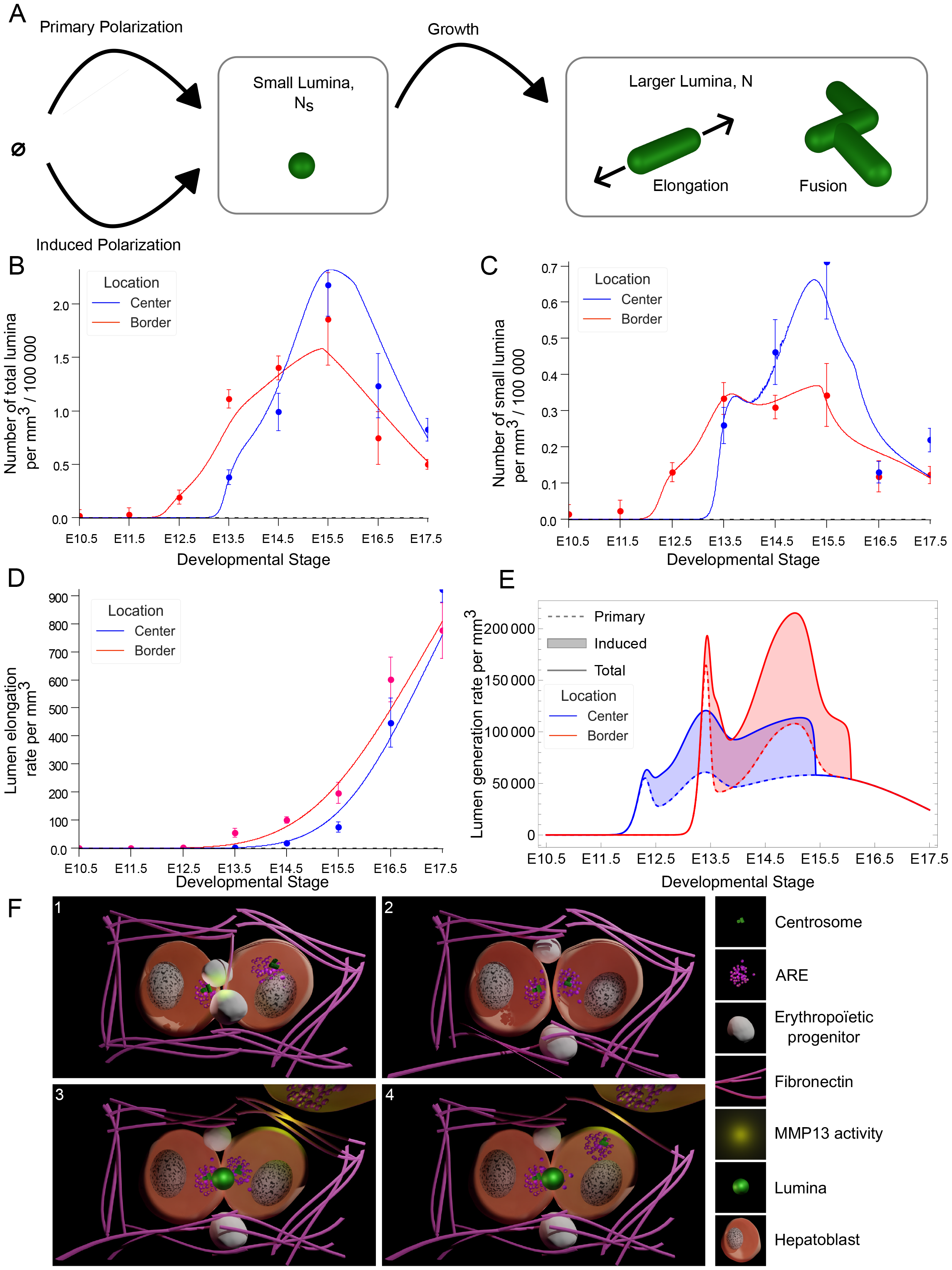
ECM degradation and lumen formation explains the observed lumen dynamics. A: Schematic of the *in silico* model. Cell division and ARE movement leads to primary polarization, ECM remodelling leads to induced polarization. Both processes contribute to the formation of small lumen. Growth leads to elongation and fusion. B-D: Comparison between the simple model and the quantification. Curves shows the model prediction Points and error bars corresponds to the data. Central area is shown in blue and the border area is shown in red. E: Prediction of the relative contribution of primary and induced polarization to the lumen generation rate measured *in vivo* at various developmental stages and in the two regions. F: Schematic of the proposed model for induced polarization.

## Discussion

Establishment of cell polarity is key to tissue morphogenesis and development. Its mechanisms have been studied extensively in different systems, both *in vivo* and *in vitro* (Datta, 2011^1^; Roignot, 2013^3^; Rodriguez-Boulan, 2014^4^; Overeem, 2015^58^; Blasky, 2015^2^). However, the mechanisms that lead to polarization in epithelial cells have been studied mostly in cells with a single apico-basal axis, and especially in MDCK cells (Datta, 2011^1^). Much less understood are the mechanisms leading to multi-axial polarization of hepatocytes (Wang and Boyer, 2004^59^; Morales-Navarrete, 2019^12^; Müsch and Arias, 2020^20^): unlike most epithelial cells, hepatoblasts establish multiple apical surfaces. This process must be coordinated both locally, to form sealed lumina between two adjacent cells, and globally, to form the three-dimensional network of bile canaliculi at the tissue level. A widely accepted model postulates that the formation of multiple lumina occurs by cell division, leading to the formation of a new lumen between daughter cells (Tanimizu, 2017^19^; Müsch and Arias 2020^20^). In this study, we show that this cell division-based model alone cannot explain the number of lumina formed during liver development. Therefore, new mechanisms are required. Here we revealed one such mechanism: hepatoblasts can generate multiple apical surfaces through repeated rounds of polarization independently of cell division. In this mechanism, after a first apical pole has been established, a population of AREs (termed pAREs) and the centrosome move across the cytoplasm to a new apical polarization site. The position of this new apical patch is dependent on ECM (specifically, Fibronectin) sensing mediated by Integrin αV. We demonstrated that the metalloprotease MMP13 is located in an intracellular pool of vesicles near an apical patch and pharmacological perturbations support its role in polarization. Altogether, our results suggest that degradation of Fibronectin by MMP13 at the apical patch of one hepatoblast signals the contacting cell to also polarize toward this site. Thus, like the cell-division based model (Tanimizu, 2017^19^), this model provides a mechanism for local coordination of placement of apical patches.

An unexpected finding of our study was that, during the polarization process of hepatoblasts, AREs near the apical surface split into two spatially distinct subpopulations. One subpopulation remained in close proximity to the initial apical surface, while the other one moved together with the centrosome to a second apical polarization site. This means that the polarization process is repeated, leading to multiaxial polarity independently of cell division. The population of AREs at the initial polarization site was retained, presumably by the MTOC, as it was previously shown that the MTOC is involved in the sub-apical retention of AREs (Le Droguen, 2015^9^; Aljiboury, 2022^60^). The association of movement of pAREs with centrosome relocation is in line with previously reported polarization events in intestinal cells (Feldman, 2012^61^). Interestingly, we observed that pARE movement is cell autonomous, i.e. not synchronized between neighboring cells. This is surprising, given that a juxtaposed cell must also polarize toward the same site to form a lumen successfully. Even more, we documented events of pARE polarization towards non-parenchymal cells *in vivo*. These ARE dynamics imply that the coordination of the apical sites of neighboring hepatoblasts is distinct from the primary polarization event. Hence, the mechanism of this coordination must direct centrosome positioning after primary polarization.

As in simple epithelial cells, the polarization of hepatoblasts depends on the sensing of the ECM at the basal site, specifically Fibronectin sensing via Integrin αV. The formation of an apical lumen depends on the asymmetric degradation of Fibronectin by MMP13. The details of how Fibronectin and its degradation by MMP13 accomplish this remain unclear. However, in addition, we suggest that Fibronectin clearance by MMP13 enables a positive feedback loop of polarization at the tissue scale: polarization leads to MMP13 secretion via centrosome positioning. This is likely accompanied with the growth of the MTOC associated with the apical surface. We propose that this growth accelerates delivery of apical components and MMP13. The latter diffuses through the tissue, leading to non-local Fibronectin degradation and hence increases polarization non-locally. Using an *in silico* model of lumen formation, elongation, and fusion, we showed that combining the mechanism of cell polarization with its positive feedback loop described here with the cell-division based mechanism can explain quantitatively the experimentally observed dynamics of lumen number and volume.

In addition to closing the tissue-scale feedback loop, the local degradation of ECM could direct the growth of neighboring canaliculi toward each other in 3D. Indeed, it was reported previously that ECM influences anisotropic growth of lumina (Li, 2016^37^). At the tissue scale this directed growth could enable lumen fusion through tight junction bridges (Figure 2D), eventually leading to a fully connected network.

The interplay between the developing bile canaliculi and sinusoidal networks could also be mediated in part by Fibronectin degradation which could provide cues for the growing sinusoidal network to avoid intersecting the growing bile canaliculi network and vice versa. In this context, it is interesting to note that, in the regenerating adult liver, endothelial cells activate stellate cells (Jarnagin, 1994^62^) by secreting a splice variant of Fibronectin normally produced by embryonic hepatoblasts (Chagraoui, 2003^57^). Upon activation, stellate cells secrete Laminin (Martinez-Hernandez, 1991^63^), which plays a role in guiding angiogenesis (Amenta, 1997^64^). Additionally, MMP13 has been implicated in the release of VEGF from the ECM and, therefore, in promoting angiogenesis (Iruela-Arispe, 2009^51^; Lederle, 2010^65^). These results suggest that the cycles of Fibronectin production and removal by hepatocytes may constitute a signal for stellate cells to guide vascularization away from the apical surfaces of hepatocytes and vice versa. Therefore, this mechanism could guarantee development of non-intersecting, intercalated bile canalicular and sinusoidal networks. Consistent with this hypothesis, it has been shown that proper differentiation of hepatoblasts is required for sinusoidal development across the liver parenchyma (Parviz, 2003^66^), potentially through VEGF signaling (Sugiyama, 2010^67^).

In summary, we propose a model for multi-axial hepatoblast polarization that relies on cell-autonomous sensing and coordination of polarization through remodeling of the local ECM environment. We also suggest this remodeling enables a positive feedback loop of polarization at the tissue scale. Extending this model to include interactions of the ECM with other cell types will lead to further understanding of the development of the architecture of the liver and its reestablishment in disease.

## Materials and Methods

### Resource availability

#### Lead Contact

Further information and requests for resources and reagents should be directed to and will be fulfilled by the lead contact, Marino Zerial.

#### Materials availability

This study did not generate new unique reagents.

### Sample preparation

#### Animal experiments

Animal experiments were conducted in accordance with German animal welfare legislation in pathogen-free conditions in the animal facility of the MPI-CBG, Dresden, Germany. Mice were maintained in a conventional barrier animal facility with a climate-controlled environment on a 12-h light/12-h dark cycle, fed ad libitum with regular rodent chow. Protocols were approved by the Institutional Animal Welfare Officer (Tierschutzbeauftragter), and necessary licenses (Licence No. DD24-5131/346/3; DD24-9168.24-9/2012-1; DD24.1-5131/451/8; DD25-5131/554/9) were obtained from the regional Ethical Commission for Animal Experimentation of Dresden, Germany (Tierversuchskommission, Landesdirektion Dresden). Mice were maintained on a C57BL/6JOlaHsd background using the internal stocks of the animal facility. The Rosa mT/mG (RRID:IMSR_JAX:007576) mouse line has already been described (Muzumdar, 2007^68^)

#### Hepatoblast Isolation

Delta-like 1 (Dlk1) positive hepatoblasts were isolated as described in (Belicova, 2021^16^), E13.5 mice were sacrificed by cervical dislocation. 12 embryonic livers were pooled and incubated in Liver perfusion media, fragmented by pipetting, and incubated for 20 min at 37[C. Liver fragments were then digested in Liver Digest Medium, supplemented with10 μg/ml DNAse I for 20 min. The suspension was then filtered through a 70 μm filter. Cells were collected by spinning at 260 g. Erythrocytes were then lysed by incubation in red blood cell lysis buffer (155 mM NH4Cl, 10 mM KHCO3, and 0.1 mM Na4 EDTA, pH 7.4) lysis was then stopped by addition of post-lysis buffer (DMEM High Glucose, 5% FBS, 2 mM L-Glutamine, 1 x NEAA). Digested cells were incubated with blocking antibody Rat Anti-Mouse CD16/CD32 (1:100) for 10 min, then with Anti-Dlk mAb-FITC (1:40) for a further 15 min. After washing with a buffer (0.5% BSA, and 2 mM EDTA in PBS), cells were incubated with Anti-FITC MicroBeads (1:10) for 15 min and separated on a magnetic column according to the manufacturer’s protocol.

#### Hepatoblast Culture

Hepatoblast culture was performed as described in (Belicova, 2021^16^). The culture wells were coated with 10 v/v % Matrigel in ice-cold PBS for at least 30 min at 37[C. Cells were then seeded in Expansion media (DMEM/F-12, GlutaMAX supplement, 10% FBS, 1× ITS-X, 0.1 μM dexamethasone, 10 mM nicotinamide, 10 ng/ml human HGF, and 10 ng/ml mouse), if no transfection was performed the expansion media was supplemented with 5 v/v % Matrigel. 24h later, the media was exchanged for Differentiation media (MCDB131, no glutamine, 5% FBS, 2 mM L-glutamine, 1× ITS-X, and 0.1 μM dexamethasone) supplemented with 5% Matrigel. Cells were seeded at a density of 30,000 cells/well in 96 well plates. If cells were transfected, they were cultured for 4 days, if not, 3 days, in both cases at 37°C, 5% CO2.

#### siRNA transfection

siRNA directed against ZO-1 and luciferase were published and validated previously (Belicova, 2021^16^). Briefly, upon plating, cells were transfected with Lipofectamine™ RNAiMAX transfection reagent (0.1 v/v%) and 10 nM siRNA. Transfection mix was removed 24h later upon addition of the differentiation media. For Integrin αV KD and associated controls, upon plating the expansion media was complemented with LNP formulated siRNA and Apolipoprotein E. Both reagents were previously published and validated (Bogorad, 2014^69^).

#### Tissue sections preparation for immunofluorescence

Pregnant mice at different stages were sacrificed by cervical dislocation. The embryos were collected, and the liver was dissected in cold phosphate buffered saline (PBS). A solution of paraformaldehyde (PFA) 4% was mixed and filtered on the day of the collection, Tween20 was added to the solution after filtration at a final concentration of 0,1%. For embryos up to E15 included the organs were placed in the PFA solution overnight at 4° on a rotating wheel. For embryos at E16 or later this step was prolonged to 2 overnight and the PFA solution was changed after the first night. The PFA was then removed, and samples were quickly washed twice in cold PBS. A third washing in PBS at 4° was then done overnight on a rotating wheel. Finally, the PBS was removed, and the samples placed in a solution of PBS + 0,05% sodium azide for storage.

The livers at E13 and later stages were embedded in 4% Agarose mixed in PBS. The septum transversum was placed at the bottom of the mold. The embedded samples were then glued on the sample holder with the septum transversum on the top side. The cutting was performed at a thickness of 100μm, an amplitude of 0.75 mm and a speed of 0,1 mm/s. Individual sections were then stored individually in PBS + 0,05% sodium azide until staining.

Sections were washed once in PBS and then permeabilized in PBS + Triton x100 0,5% for one hour. The sections were then incubated with primary antibodies in a solution of PBS+ Triton x100 0,3%+ Fish gelatin 0,2% for two days at room temperature. Sections were then washed 5 times 15 minutes in PBS+ Triton x100 0,2% before starting the incubation with secondary antibodies (in the same buffer as primary antibodies) for two days at room temperature. The sections were then washed 5 times 15 minutes in PBS+ Triton x100 0,2% and 3 times 5 minutes in PBS. The sections were then further processed through the clearing protocol.

The clearing was performed following the SeeDB protocol (Ke, 2013^70^). Sections were incubated in increasing concentration of fructose on a rotating wheel at room temperature. Samples were placed in fructose 25% in distilled water for 6h then fructose 40% for 6 hours then fructose 60% overnight then fructose 80% overnight then fructose 100% overnight. Finally, samples were incubated in a See Deep Brain (SeeDB) solution overnight and then mounted in the same solution.

#### Tissue sections preparation for EM

Liver was dissected from E13.5 mouse embryos and cut into a few pieces, which were immersion fixed in 4% PFA 200 mM HEPES, pH 7.4 for 24 hours. The tissue was then washed with dH2O, the residual fixative quenched with 0.1% glycine in dH2O before sectioning. All sections were analysed in a Tecnai T12 transmission electron microscope, operated at 100 kV and equipped with an axial 2k CCD camera.

The pieces were embedded in 12% bovine gelatine and infiltrated with 2.3 M sucrose in dH2O on a rotating wheel at room temperature for at least 24 hours. Such cryo-protected pieces were mounted on pins and snap frozen in liquid nitrogen to become hard. 90-nm thin Tokuyasu thawed cryo-sections were prepared using a Leica FC6 cryo-ultramicrotome. Sections were transferred to formvar-coated, 100 mesh, hexagonal, copper grids. Grids were sequentially incubated on drops with first 1% glycine in PBS for 2 min, then 1% cold water fish skin gelatine in PBS for 1 minute, then Aminopeptidase N/CD13 antibody (1:100) for 30 minutes, then unconjugated bridging rabbit anti-rat IgG (1:500) for 15 min, finally 10-nm protein A gold (1:25) for 30 min.

Labelling reagents were diluted in 1% cold water fish skin gelatine in PBS, and PBS was used for washing after incubation with the labelling probes. After immunolabeling, grids were washed with dH2O and embedded in a thin layer of 0.2% uranyl acetate in 1.8% methyl cellulose prepared in dH2O.

Imaging was then performed using either an Axio Examiner.Z1 (Zeiss) or an Axio Observer (Zeiss) microscope with a Zeiss LCI Plan-Neofluar 63x 1.3 Gly/Water DIC objective.

#### Primary cells preparation

Cells were fixed overnight at 4° at different times after plating, using 4% PFA in PBS buffer with 0,1% Tween20. Cells were then permeabilized with a solution of PBS + Tritonx100 at 0,3%. Then cells were blocked using a solution PBS + Tritonx100 0,1% + Bovine serum albumin (BSA) 3% for 2h at room temperature. Cells were then incubated overnight with primary antibodies in the blocking buffer. Cells were then washed 3 times 5 minutes with PBS + tritonx100 0,1%. Then cells were incubated overnight with secondary antibodies in the blocking buffer. Finally, cells were washed 3 times with PBS and imaged within a few days.

Secondary antibodies were used at the concentration of 1/1000. For actin staining we used Phalloidin labelled with Alexa 488 at a concentration of 1/200. Dapi was used at a concentration of 1/5000.

Imaging was then performed on an Olympus – IX83 microscope with an Olympus UPLSAPO objective.

### Image analysis

#### Liver sections visualization and quantifications

Tissue section quantifications were performed using the Imaris software (Oxford Instruments).

For bile canaliculi we generated a colocalization channel based on CD13 and phalloidin. We then placed a threshold to capture most of the objects. At this point several ARE were still present in the pool. To remove them we sorted out the objects by ZO-1 intensity. Based on the histogram of the average intensity it was clear which ones were positives and which ones were negatives.

For apical recycling endosomes we generated a colocalization channel based on CD13 and Rab11. We then placed a threshold to capture most of the objects. At this point several lumina were still present in the pool. To remove them we sorted out the objects by phalloidin intensity. Given that lumina display a very strong phalloidin staining it was clear based on the signal intensity histogram which ones were lumina and which ones were ARE.

#### Hepatoblast culture quantifications

The analysis of bile canaliculi morphology and bile canaliculi network properties was performed using a previously published script (Mayer, 2023^71^) running on Fiji Fiji (Schindelin, 2012^72^). Briefly, Images were imported using Bioformats (Linkert, 2010^73^) and the aspecific staining (appearing as extracellular puncta of saturated pixels) was segmented using a simple threshold combined with an inflation and then removed from the images. The Apical signal and the actin signal were smoothened with a median filter and segmented independently with the Huang (Huang & Wang, 1995^74^) and (Doyle, 1962^75^) methods respectively. The overlap between the two segmented images was then kept as the basis to identify apical patches. The apical patches were then completed by a series of inflations followed by the “fill holes” function and then a series of deflations with morpholibJ (Legland, 2016^76^). At this point a size threshold is applied on the remaining potential lumina and the lumina touching the borders of the images are removed also with morpholibJ. A background removal is then applied on the junction channel, a median filter is applied followed by a segmentation with the default method for the autothreshold Fiji plugin. A size filter is applied, and the remaining segmented junctions are used as seeds for a geodesic reconstruction (using morpholibJ) performed onto the potential lumina. The features are then exported as csv and plotted in python using matplotlib and seaborn.

Statistical analysis was performed using an unpaired Student’s two-tailed two-sample t test of the means of each biological replicate using python.

Live images were restored using Noise to void (Krull, 2020^77^).

#### Motion Tracking

The relative spatial distribution quantification was performed on MotionTracking for its versatility at the analysis level.

We corrected for channel misalignment using the phalloidin channel as a reference and then applied several gaussian blur filters to the images. We then segmented the CD13 channel and the basal cue candidate channel using an approach similar to (Morales-Navarrete, 2015^78^).

## Author contributions

J.D.: Conceptualization, Investigation, Formal Analysis, Methodology, Visualization, Writing – original draft; J.I.V.: Conceptualization, Investigation, Writing – review & editing; M.Bo.: Conceptualization, Investigation, Methodology, Visualization, Writing – original draft; N.P.M.: Conceptualization, Software, Visualization; L.B.: Conceptualization, Investigation, Writing – review & editing; U.R.: Investigation; M.Be.: Investigation, Writing – review & editing; S.S.: Investigation, Resources; P.A.H.: Methodology, Conceptualization, Writing – review & editing; Y.L.K.: Methodology, Conceptualization, Software, Writing – review & editing; M.Z.: Conceptualization, Supervision, Writing – review & editing, Funding acquisition

## Acknowledgements

We are grateful to Timofei Zatsepin (Skoltech, Moscow) for providing the siRNA against ZO-1 and Anylum pharmaceutical for providing the LNP formulated siRNA against aV.

We are also grateful to Drs Wieland B Huttner, Elisabeth Knust, Ivo Sbalzarini, Frank Jülicher, Quentin Vagne, Aleksandra Sljukic, Aparajita Lahree, Lisa Johnsen and Carlotta Mayer for valuable discussions.

We would like to acknowledge Jan Peychl, Sebastian Bundschuh and Britta Schroth-Diezfrom the light microscopy facility; Katrin Reppe and Anke Muench-Wuttke from the biomedical services facility at Max Planck Institute of Molecular Cell Biology and Genetics in Dresden for their contributions.

This research was financially supported by the German Federal Ministry of Research and Education (BMBF, SysBioII, grant number 031L0044), the European Research Council (ERC) under the European Union’s Horizon 2020 research and innovation program (RULLIVER, grant no. 695646), and the Max Planck Society. We would like to acknowledge the Center for Information Services and High-Performance Computing (ZIH) at the Technical University of Dresden for the generous allocation of computer time.

## Appendix

### 1 Model description

We experimentally observe the number of bile canaliculi lumina and their volume over the course of development. Additionally, we partition small lumina with volumes less than 10 μm^3^, and track their number as well. Here we construct a compartmental model of the dynamics of these quantities by modeling four processes: generation of new lumina, maturation of small lumina into elongating ones, elongation of those lumina, and fusion of lumina. These processes occur in a growing tissue, which we also account for in our model.

We seek differential equations for the dynamics of each of the three experimental variables: number of small lumina, number of total lumina, and total volume. In this section we first describe our model for the growth of the tissue. Then we detail our models for rates of each of the processes. Finally, we assemble these rates and state the differential equations for the full model.

#### 1.1 Growth of the liver

The liver grows significantly over the course of our measurements, which we seek to account for in our model. Both liver volume and hepatocyte number have been measured previously. We use the data of Paul et al. [1]. This approach is similar to that of Weiss et al. [2]. Paul et al. [1] measure liver weight, which we transform to volume by approximating the density as that of water. We note the liver likely has a density different than that of water and that its density likely changes over the course of development due to the changing composition of cells and ECM. However, we find it probable that both the overall difference in density between the liver and water and changes in density over development are small compared the density of water. Additionally, such differences are unlikely to qualitatively affect the outcome of our model, which aims to capture general mechanisms only. We find the volume density of hepatocytes by dividing the measured total hepatocyte number by the volume. We fit continuous functions to these data to arrive at equations approximating total liver volume Ω(*t*) and volume density of helpatocytes *c*(*t*),

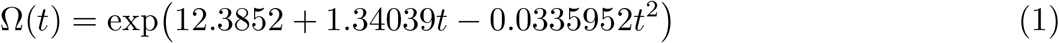

and

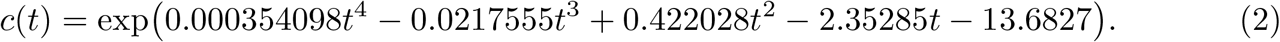

**Figure 1:**
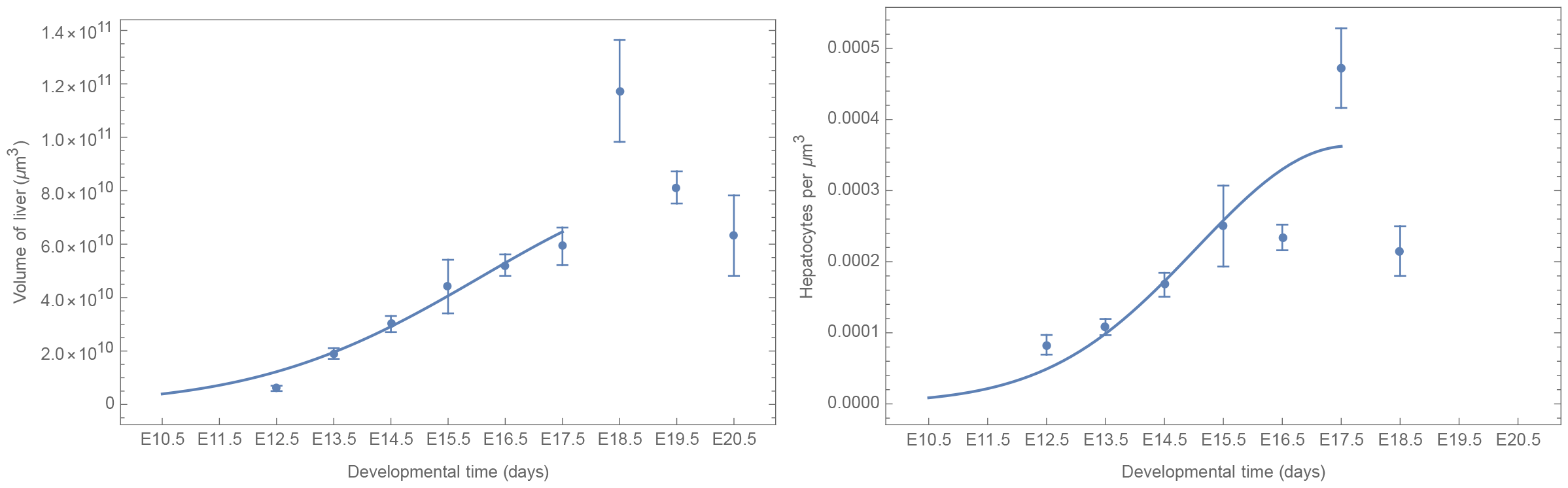
Data from Paul et al. [1] and fit functions for liver volume Ω(*t*) and hepatocyte volume density *c*(*t*).

We show these fits together with the transformed data from ref. [1] in figure 1.

#### 1.2 Generation of new lumina

New lumina are generated by polarization of hepatocytes. We include several polarization mechanisms into the model. We consider polarization by cell division as well as two cell division independent mechanisms.

##### 1.2.1 Cell-division dependent polarization

We estimate the rate of hepatocyte divisions by taking the derivative of our fit to the data from ref. [1]. The volume density of cell divisions in the liver is the derivative of the total number of divisions divided by the volume, i.e.

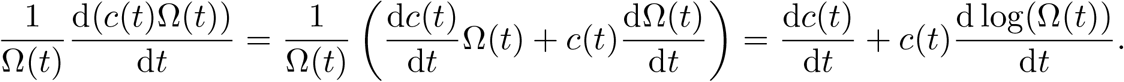

Then, we state the rate of cell division dependent generation as

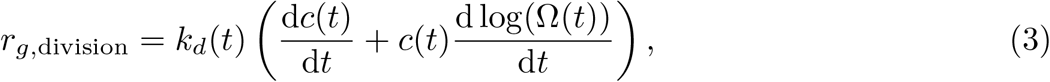

where we have included the step-like function

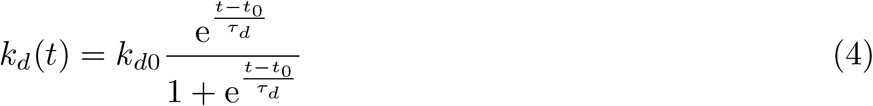

to allow cell divisions to begin generating lumina at some time during development. We imagine a developmental signal determines the turn on time. The fit parameters *t*_0_ and *τ*_*d*_ determine when and how quickly this happens, respectively. If *τ*_*d*_ is small and *t*_0_ goes to early times, this function becomes effectively constant at *k*_*d*0_. We note *k*_*d*0_ should be between 0 and 1, as we expect one cell division event to create at most one lumen.

##### 1.2.2 Cell-division independent polarization

Our experiments reveal that hepatocytes polarize independently of cell division, mediated by repositioning of the apical recycling endosome as described in the main text. We model two kinds of cell division independent polarization. The first is polarization induced by ECM degradation. The second is polarization which is not dependent on cell division, nor induced by ECM degradation. We imagine this happens in response to the same external cue that turns on cell division dependent polarization. We term this second type of polarization spontaneous.

###### 1.2.2.1 Induced polarization

As discussed in the main text, we observe that hepatocytes which have already polarized can reposition the apical recycling endosome near areas of low ECM to generate another new lumen. Since hepatocytes continue to degrade the surrounding ECM as they generate more lumen volume over the course of development, we suppose these mechanisms interact to form a feedback: as more volume is formed, more ECM is degraded, leading to more lumen formation and so on. As such, we model this induced polarization to be proportional to the current lumen volume density and the number density of new lumina, with rate

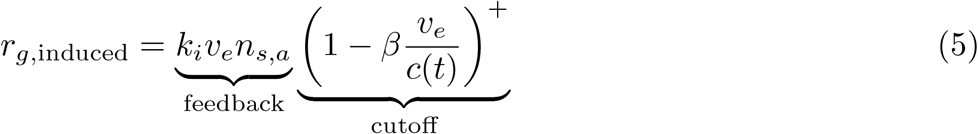

where the rate constant *k*_*i*_ is a fit parameter, *v*_*e*_ is the volume of elongating lumina, and *n*_*s,a*_ is the number density of small lumina which are able to induce generation. We expand on the construction of *n*_*s,a*_ in the following section.

This feedback loop is cut off as hepatocyte surfaces become saturated with the maximum amount of apical surface they can support. We therefore include a term that cuts this generation off as lumen volume per cell increases. The lumen volume density at which this occurs is determined by the fit parameter *β*. We note we take only the positive part of this function, denoted by ^+^, so that the induced generation rate is 0 once the critical volume is exceeded.

###### 1.2.2.2 Spontaneous polarization

We suppose spontaneous generation is proportional to the number of cells, but with a rate that has a peak in time, created by some external signal. We state the rate of spontaneous generation as

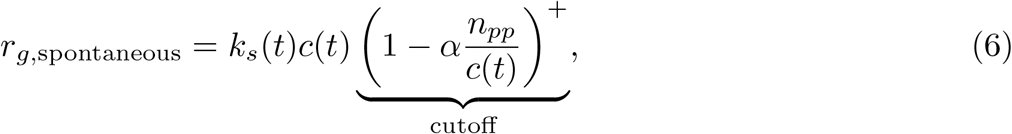

where the peaked function

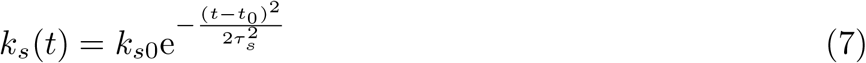

has fitting parameters *k*_*s*0_, *t*_0_ and *τ*_*s*_ which determine the height, peak time, and width, respectively. With large values of *τ*_*s*_, this function approaches being constant in time. We note this *t*_0_ is the same that appears in equation 4, which defines the turn on time of cell division dependent lumen generation. Our minimal model is that one signal is responsible for both.

We suppose hepatocytes can only generate a limited number of primary lumina, where primary lumina are those not generated by induction. The reasoning for this is expanded on in section 1.6.

Therefore, we introduce a term which cuts off spontaneous generation as the number of primary lumina (*n*_*pp*_) per cell increases. The fit parameter *α* determines the maximum number of lumina generated by primary polarization per cell.

#### 1.3 Maturation of small lumina into elongating ones

We observe that lumina are approximately spherical when their volumes are below 10 μm^3^ (main text figure 1). This suggests that lumina grow spherically up to this volume, then begin to elongate. We term growth of a lumen between being generated and exceeding 10 μm^3^ maturation, and model it as taking some characteristic time *τ*_mat_, which is a constant over development. This translates to a rate of maturation

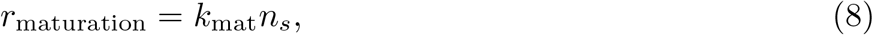

where *n*_*s*_ is the number density of small lumina and *k*_mat_ = 1*/τ*_mat_ is the rate constant.

#### 1.4 Elongation of lumina

We observe that non-small lumina have roughly the same radius, regardless of their length. This suggests that after a small lumen exceeds 10 μm^3^, it begins to elongate roughly as a cylinder of constant radius. Elongation happens from both ends, so a new elongating lumen has two growing tips. Larger elongating lumina typically have more than two growing tips, with the extra tips having arisen from fusion events or potentially branching. Our simple model for these dynamics is that the number of growing tips scales with the volume of a lumen roughly as the surface area of a characteristic sphere, or with the 2*/*3 power of volume. Then the elongation rate is proportional to the surface area of the average lumen and the number density of elongating lumina,

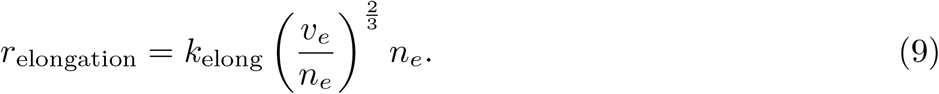

Here *k*_elong_ is the rate constant of elongation.

#### 1.5 Fusion of lumina

The process by which elongating lumina encounter each other and join is not well understood. In general, it is a complex spatial process which must depend on the distance between lumina, their size and the geometry of the hepatocyte surfaces in between, as well as any mechanisms hepatocytes might have to guide the elongation toward other lumina. We do not attempt to resolve these complexities in this simple model. A more in depth understanding of lumen fusion could help make this term more accurate, but requires significant analysis. While of great interest, a more detailed model of fusion would likely put the precision of this term beyond the precision of this model overall and not aid its explanatory value.

As stated in the previous section, lumina grow at their tips. One might then expect a fusion term proportional to 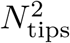, if fusion was reciprocal, or *N*_tips,1_*N*_tips,2_ for two subpopulations which fuse asymmetrically. We assume most fusion occurs between a few, large lumina and many smaller but still elongating lumina. The few large lumina have a number of tips determined by the surface of a characteristic sphere, as discussed in the previous section. Smaller elongating lumina have two tips. This yields *N*_tips,1_ *≈* (*V*_large_*/N*_large_)^2*/*3^, *N*_tips,2_ *≈* 2*N*_*e*_. We further assume something to get *V* only. Then the rate of fusion is

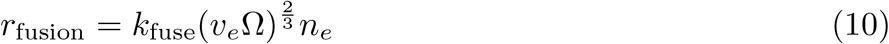

where we have absorbed the factor of 2 into the rate constant *k*_fuse_. This simplified model is more accurate later in development, when the majority of fusion is occurring. It is not a good approximation for fusion between lumina of similar sizes at early times, but such fusion events are rarer.

#### 1.6 Differential equations

With the rates of the processes we wish to model in hand, we proceed to construct the full model. We note that we track the per liver volume density of lumen number and lumen volume, which necessitates the inclusion of a term tracking the density change with change of volume of the liver. This term has the form 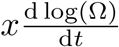 for a density quantity *x*. To see the origin of this term, examine the derivative of the density in terms of the total quantity *X* = *x*Ω:

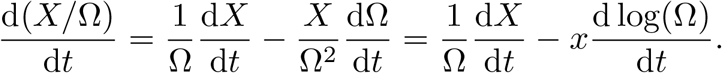

We track several more refined compartments, then reconstruct the model variables. We first distinguish induced polarization from the other two types. Together, we term lumina produced either by cell division or spontaneous generation as primary. The rates of primary polarization and induced polarization are then

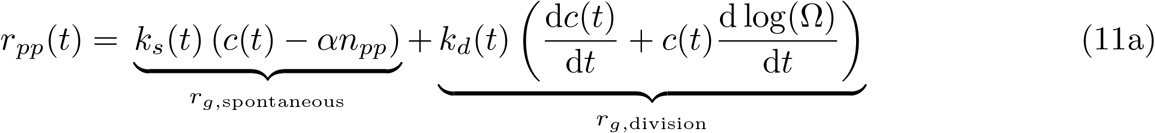

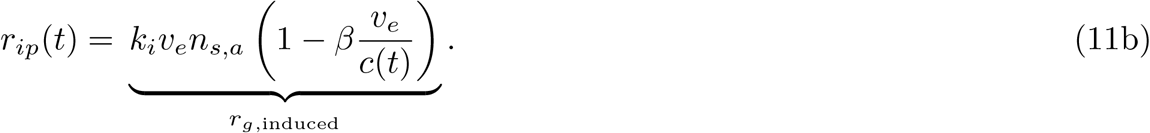

This allows us to track the total number of lumina produced by primary polarization, *n*_*pp*_, and limit their number in the model. We also limit the ways induced lumina can be produced: first, an induced lumen is not able to induce more lumina. Second, each primary lumen is able to induce only one more lumen. Therefore we split the number of small lumina into two compartments, those which are able to induce polarization, *n*_*s,a*_, and those which are unable to do so, *n*_*s,u*_. When a primary lumen induces generation, it is transferred from the first compartment to the second.

We also split the volume of elongating lumina *v*_*e*_ from the total volume and track it. It grows by elongation and entry of the volume of small lumina into the elongating compartment.

With this, the differential equations describing the dynamical variables are

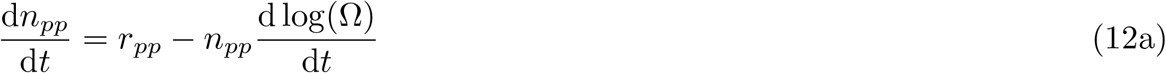

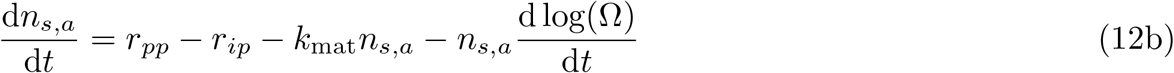

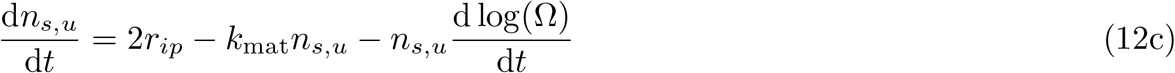

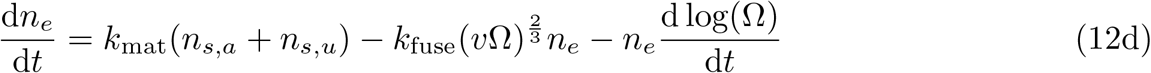

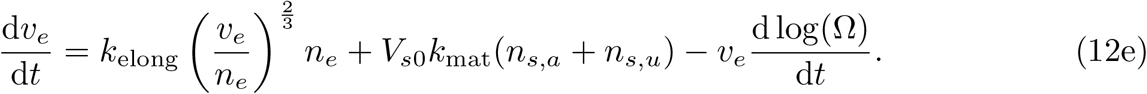

From these dynamical variables we reconstruct the model variables, the number of small lumina, *n*_*s*_, the total number of lumina, *n*, and the total volume of lumina, *v*, all measured per liver volume:

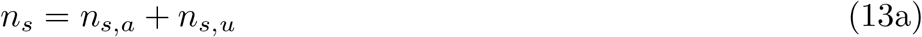

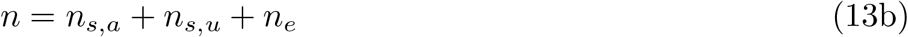

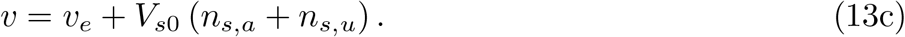

These variables are the ones we compare with the experimental data.

##### Cell division only model

We note that if we set the rates of cell division independent generation to zero, the model simplifies significantly to

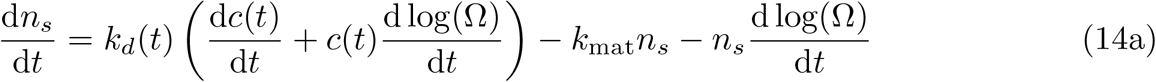

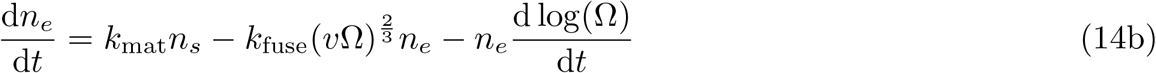

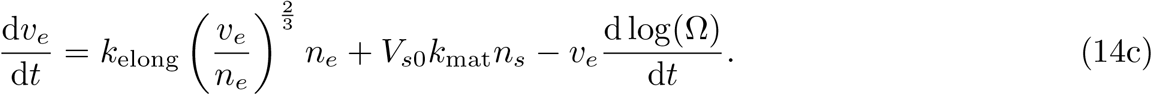

The model variables are then

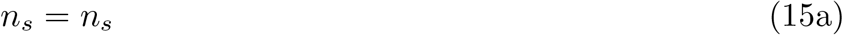

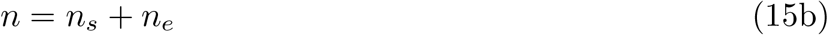

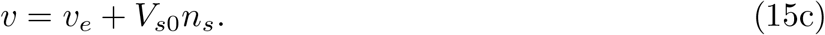

### 2 Model fitting

Sets of equations for the center and border were fit simultaneously to experimental data on number of small lumina, number of large lumina, and total volume. The fit was accomplished using the Fit Model software [3] (http://pluk.mpi-cbg.de/projects/fitmodel), which performs a *χ*^2^ minimum search. The best fit parameters for the full model are given in Table 1. All parameters are global save the two which determine the amplitude and timing of the initial polarization pulse (*k*_*d*0_ and *t*_0_), which take on separate values for the center and border. The initial conditions for all model variables were 0 at day E10.5.

**Table 1:** Fitted parameter values for full model. Lengths, areas and volumes refer to measurements of lumina. d: days.

| Parameter | Value | Units | ± 95% CI |
| --- | --- | --- | --- |
| Global |  |  |  |
| $V_{s0}$ | 4.07 | $\mu\text{m}^3$ | $1.7 \times 10^{-2}$ |
| $\alpha$ | 2.91 | cells per lumen | $1.7 \times 10^{-4}$ |
| $\beta$ | $1.17 \times 10^{-3}$ | cells per $\mu\text{m}^3$ | $2.68 \times 10^{-5}$ |
| $k_{d0}$ | 0.36 | lumina per cell | $3.13 \times 10^{-3}$ |
| $k_{\text{mat}}$ | 2.67 | $\text{d}^{-1}$ | $5.8 \times 10^{-4}$ |
| $k_{\text{elong}}$ | 6.97 | $\mu\text{m d}^{-1}$ | $1.5 \times 10^{-3}$ |
| $k_{\text{fuse}}$ | $4.83 \times 10^{-7}$ | $\mu\text{m}^{-2} \text{d}^{-1}$ | $5.0 \times 10^{-9}$ |
| $k_i$ | $6.60 \times 10^4$ | $\text{d}^{-1}$ | $5 \times 10^{-1}$ |
| $\tau_d$ | 0.20 | d | $3 \times 10^{-6}$ |
| $\tau_s$ | 0.29 | d | $1.2 \times 10^{-5}$ |
| Center region |  |  |  |
| $k_{s0,c}$ | 32.3 | lumina per cell $\text{d}^{-1}$ | $8.5 \times 10^{-3}$ |
| $t_{0,c}$ | 12.4 | d | $1 \times 10^{-4}$ |
| Border region |  |  |  |
| $k_{s0,b}$ | $1.17 \times 10^3$ | lumina per cell $\text{d}^{-1}$ | $1.7 \times 10^{-1}$ |
| $t_{0,b}$ | $t_{0,c}+1.46$ | d | $7 \times 10^{-5}$ |

Before E13.5, the liver is small and cannot be differentiated into center and border regions. Imaging volumes are centered on the hilum, but cover most of the organ. It is not immediately clear, therefore, how these data points should be included in the fits. We choose to include the data points in the fits for the center, but not the border. Our reasoning for this is as follows: the imaging volumes at days E10.5–E12.5 certainly contain the tissue that will become the center, so the data represent our best guess for the lumen numbers and volume in the center at these time points. While the imaging volumes also contain some tissue that will become the border region, we find it likely that the mechanism which causes the differences we observe at later time points is mediated by a signal generated at the center of the organ. Under such a model, the length scale of this signal would determine where center transitions into border. Since the liver is small at days E10.5–12.5, we reason that most or all of the organ may below this length scale and behave as center at these early times. Therefore we choose to leave the fit for the border unconstrained before E13.5, at which time a clear difference between center and border is observed.

### 3 Model detailed results

The results of the model for the number of small and total lumina, as well as for total volume are shown in main figure 9. We replot them here in figure 2. We expand here on the model results, with a focus on the number of lumina generated by the different mechanisms.

**Figure 2:**
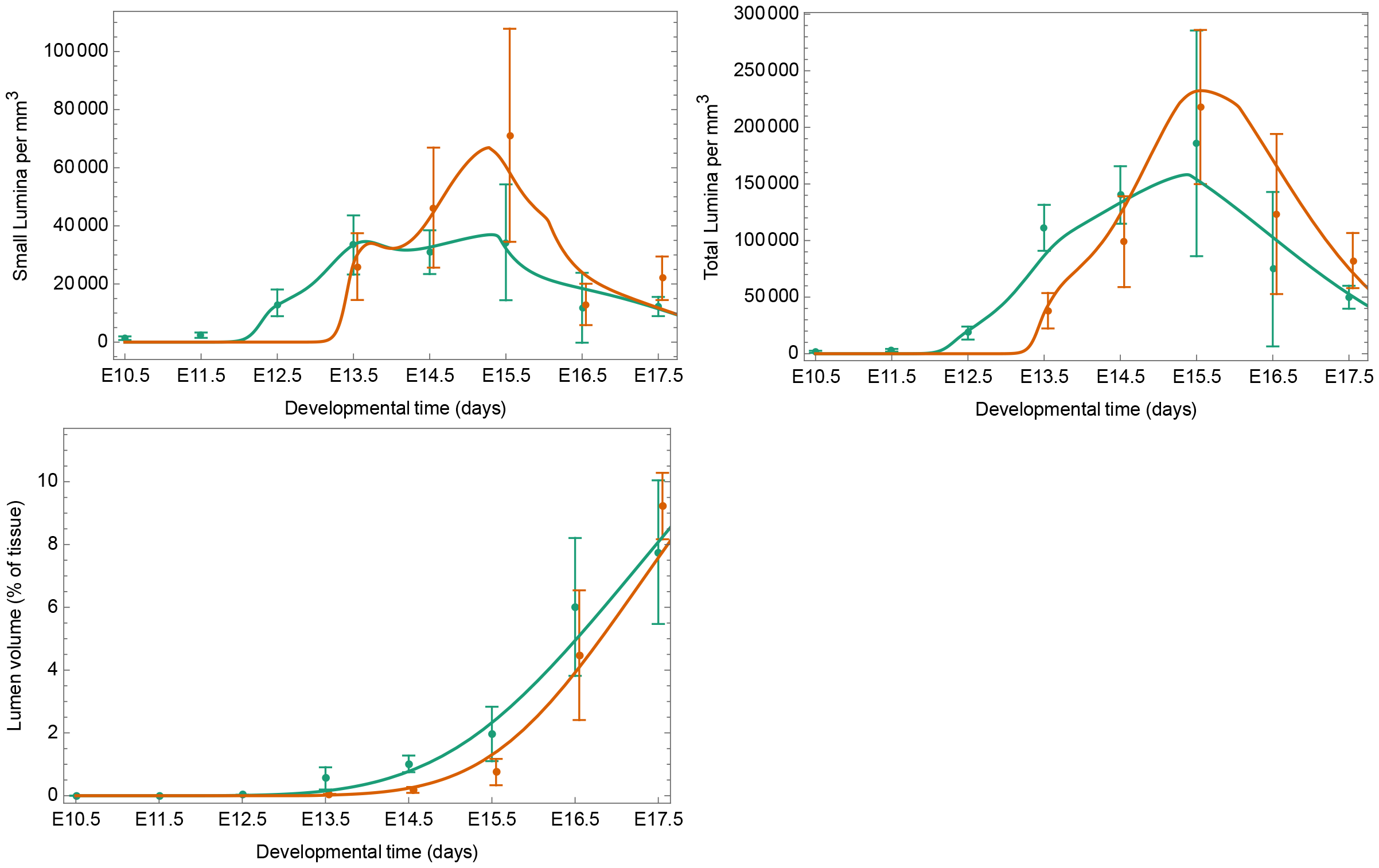
Results of the full model Results for the center are shown in green, those for the border are shown in orange. Experimental values are shown as mean*±*95% confidence interval. Values for the border are shown slightly shifted for visual clarity.

With the fitted model we can compare each of the generation rates over the course of development. Each of the three rates are shown for the center and border regions in figure 3. In the center, we find that spontaneous polarization is high early, followed by peaks of induced and cell division dependent polarization. By E15.5, the model predicts that most hepatocytes are saturated with lumina, so new generation thereafter is dependent on generation of new hepatocytes by cell division. In the border region, the model predicts similar dynamics, but starting later during development and with higher instantaneous rates. As in the center region, cell division dependent polarization dominates after E16.5. The model predicts the two regions have different fractions of lumina generated by each mechanism. In the center, about half are generated by cell division dependent polarization, with 1/8 generated spontaneously and the other 3/8 generated by induction. In the border, about 1/3 of lumina are generated by cell division, while about 1/4 are generated spontaneously and the remaining 4/10 are induced.

**Figure 3:**
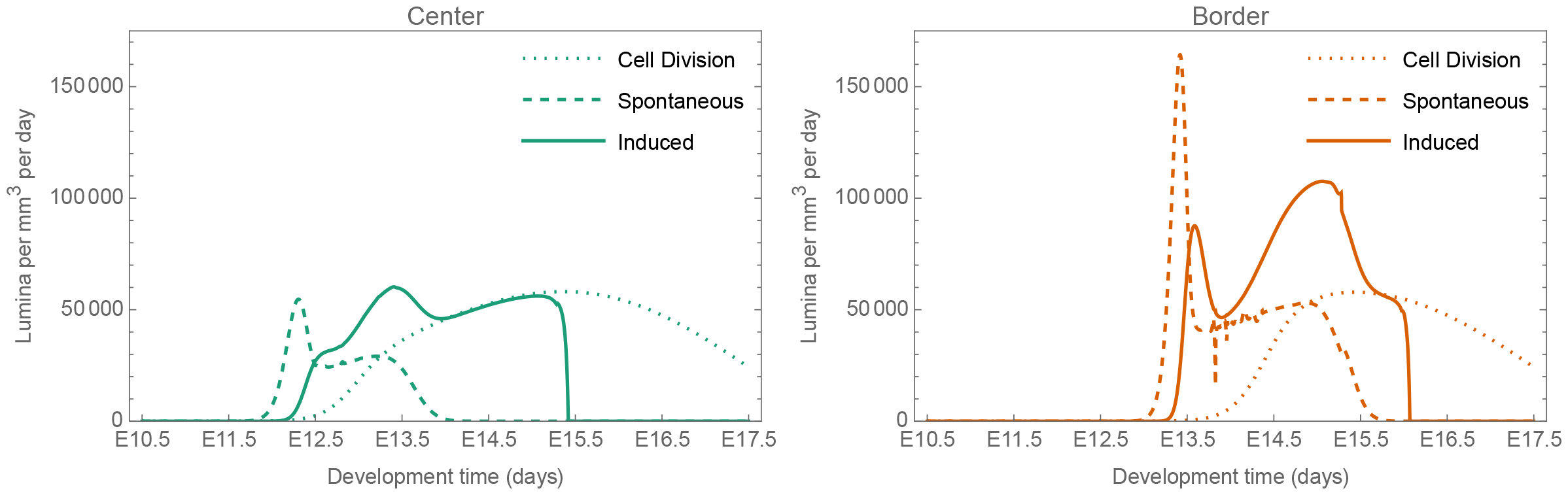
Generation rates of new lumina from each of the three mechanisms over the course of development. Rates for the center are shown in green, rates for the border are shown in orange.

### 4 Comparison to alternative models of lumina generation

In this section we investigate the contributions of each mechanism to the fit of the model to the data by comparing versions of the model with corresponding terms removed.

#### 4.1 Model with cell division mediated polarization only

When we remove both types of cell division independent polarization from the model and refit the parameters, we find that fits are not able to reproduce important qualitative features of the data. To give the most generous possible interpretation to the cell division only model, we constrain the fit to give exactly one lumen per cell division. This number is likely an overestimate as it would predict some hepatocytes with 5 or more lumina (hepatocyte number increases from .5 to 30 million from E12.5 to E18.5 [1], and .5*∗* 2^*n*^ = 30 gives *n* = 5.8 divisions). The parameters resulting from this fit are given in table 2. Despite the high value for lumina produced per cell division, this model significantly underestimates the number of small lumina at the border, as it is able to account for only 65% of the lumina observed. Fits are shown in figure 4. While the number of parameters to fit is lower, the fit of this model is also worse on balance with a BIC value of 335, compared to 217 for the full model.

**Table 2:** Fitted parameter values for the cell division only model. Lengths, areas and volumes refer to measurements of lumina. d: days.

| Parameter | Value | Units | ± 95% CI |
| --- | --- | --- | --- |
| Global |  |  |  |
| $V_{s0}$ | 1 | $\mu\text{m}^3$ | 1.45 |
| $\beta$ | $2.58 \times 10^{-2}$ | cells per $\mu\text{m}^3$ | $<1 \times 10^{-7}$ |
| $k_{d0}$ | 1 | lumina per cell | |
| $k_{\text{mat}}$ | 4.48 | $\text{d}^{-1}$ | 0.2 |
| $k_{\text{elong}}$ | 6.91 | $\mu\text{m d}^{-1}$ | $1.94 \times 10^{-1}$ |
| $k_{\text{fuse}}$ | $8.61 \times 10^{-7}$ | $\mu\text{m}^{-2} \text{d}^{-1}$ | $5.23 \times 10^{-8}$ |
| $\tau_d$ | 0.25 | d | $8.64 \times 10^{-2}$ |
| Center region |  |  |  |
| $t_{0,c}$ | 11.2 | d | $1.84 \times 10^{-1}$ |
| Border region |  |  |  |
| $t_{0,b}$ | $t_{0,c}+1.37$ | d | $1.44 \times 10^{-1}$ |

#### 4.2 Comparison to models with mechanisms removed

The mechanisms we propose in the full model allow it to find a good fit to the data. However, since fit has many parameters, it should be expected to perform well. To understand how much the additional mechanisms improve the fit of the model, we fit several nested models in which we remove some of the mechanisms. In figure 5, we show the results of 3 additional models: one in which we remove induced polarization, one in which we remove spontaneous polarization, and one in which we allow only spontaneous polarization. We find that each of these additional models are able to fit the general dynamics of the data. However, each suffers from not being able to fit the data well in some region. For example, the model with spontaneous generation only is unable to fit the number of small lumina at late times. Quantitatively, the full model has both the best fit and the best BIC value, indicating that the additional complexity introduced by each of the mechanisms is a worthwhile addition to the model. *χ*^2^ and BIC values for each model are shown in table 3.

**Table 3:** Statistics for the different models with mechanisms removed.

| Model | $\chi^2$ | BIC |
| --- | --- | --- |
| Full | 80.953 | 217.223 |
| No induced | 99.5837 | 246.582 |
| No spontaneous | 99.7317 | 238.976 |
| Only spontaneous | 112.512 | 264.536 |

**Figure 4:**
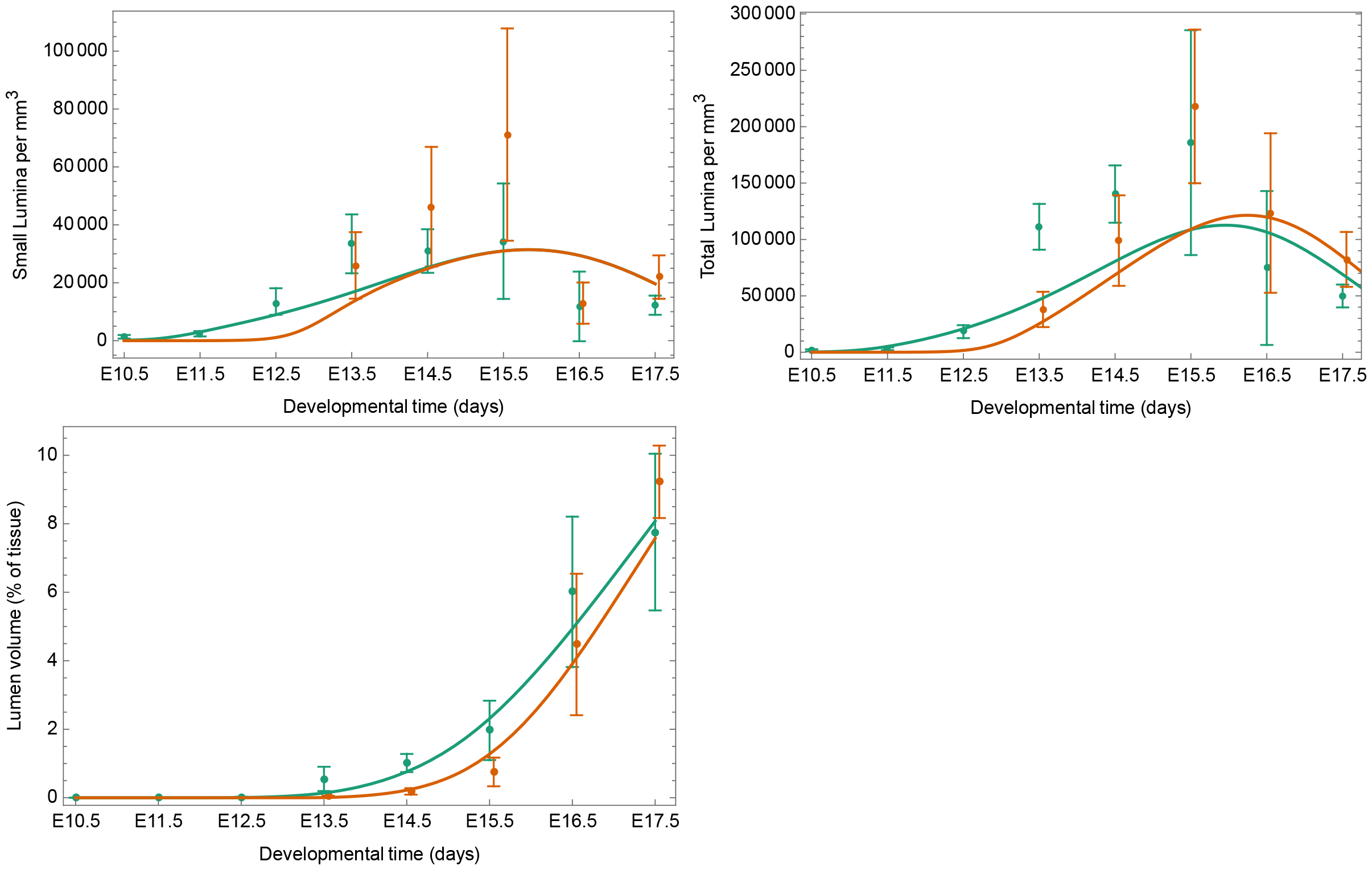
Model results with cell division mediated generation only. Results for the center are shown in green, those for the border are shown in orange (shifted slightly to aid visualization).

**Figure 5:**
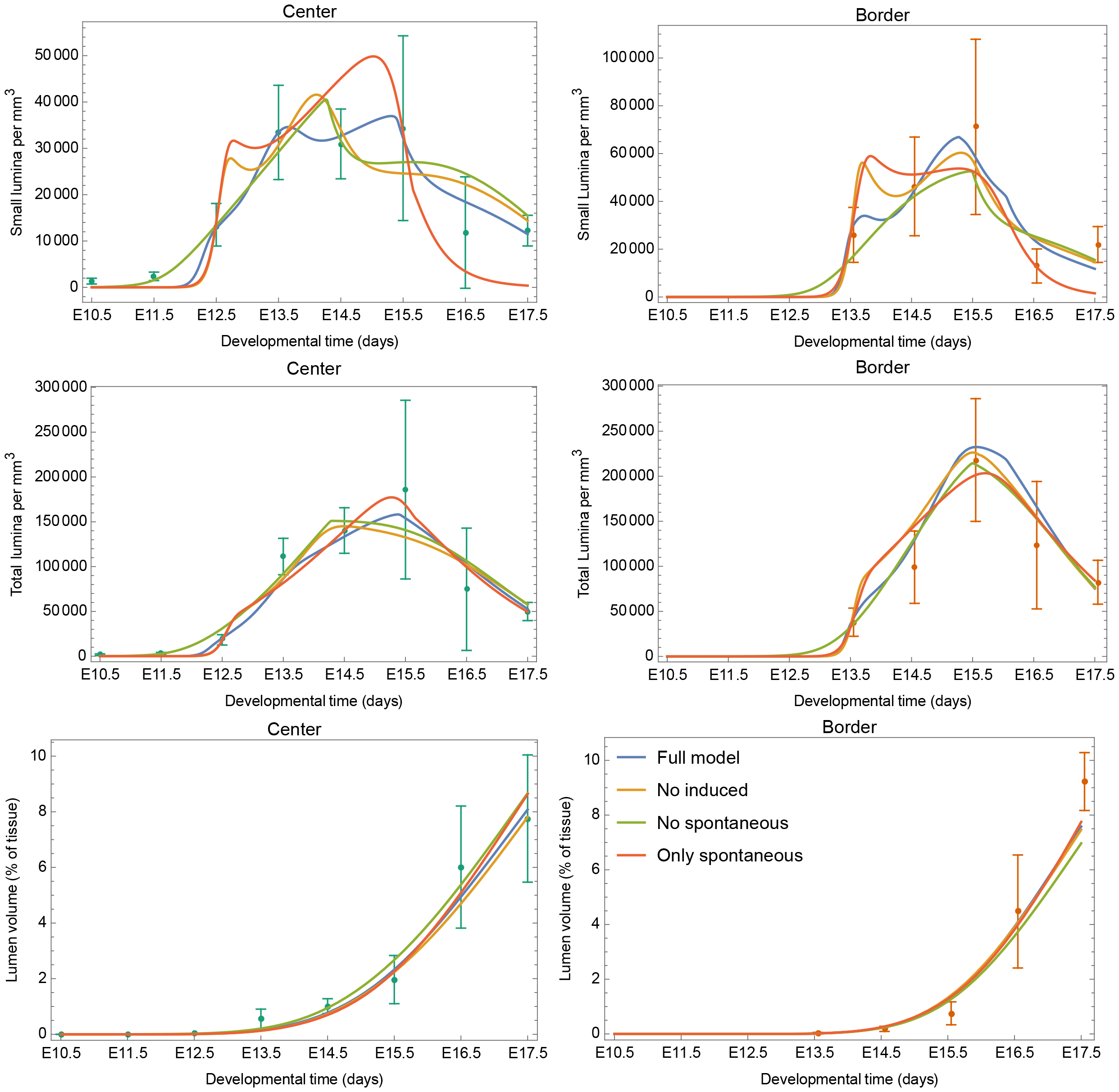
Results for the full model and the three models with mechanisms removed Colors of curves in each panel correspond to the legend in the bottom right.

---

**Supplementary Figure1: Differences across the embryonic liver tissue**.

A: Illustration of the liver section plane on an E14.5 embryo and maximum projection over 100 μm of a whole liver section immunostained against an apical surface marker (CD13). ST: Septum Transversum, H: Hilum. Scale bar: 1mm

B: Maximum projections of two region: close to the border (border) and close to the hilum (center) over 30 μm of an E14.5 embryonic liver section, immunostained against an apical surface marker (CD13). Same sample as A. Scale bar: 20μm

**Supplementary Figure2: ARE and lumen morphology**.

A: 3D reconstruction of lumina (CD13+, ZO-1+, F-actin+) and ARE (CD13+, ZO-1-, F-actin-) of liver sections, in the central region of the organ, between the initial lumen formation and network formation. The white box shows the image frame, ticks are 10μm.

B: ARE quantifications across the considered timeframe. Quantifications from 3 sections per timepoint imaged at the center and border.

C: Single plane image of an immunostaining directed against luminal markers (CD13, ZO-1) and a cell border marker (F-actin labelled with Phalloidin), on fixed liver section at E14.5 in the hilum region. Dashed white square shows the region selected for single channel magnifications. Green arrowheads show intracellular CD13 (ARE). Red arrowheads show apical membrane initiation site: narrow CD13+/ZO-1+/F-actin+ with ARE spanning towards the inside of the cell. Red Stars show fully formed lumen: large CD13+/ZO-1+/F-actin+ structure. Scale bar: 10μm

**Supplementary Figure3: ZO-1 KD effect on lumen morphology**.

A: Single plane images of immunostaining directed against ARE markers (CD13, Rab11) and a cell border marker (F-actin labelled with Phalloidin), on fixed primary hepatoblasts, without treatment (NT) or after treatment with siRNA directed against Luciferase (non-targeting - Luc) or ZO-1. Scale bar: 10μm

B: Effect of ZO-1 KD on lumen morphology. 20 images per well, 2 wells per replicate, 2 replicates per condition. p-values are shown when lower than 0.05.

**Supplementary Figure4: Basal markers distribution**

Single plane images of E14 liver sections immunostained against various basal markers and CD13. Scale bars: 10μm

**Supplementary Figure5: MMP inhibitor screen**

Table showing the targets of the used mmp inhibitors (dark cases) and the observed output (lumen formation observed or not).

