## Supplementary figures and images for "Hepatoblast iterative apicobasal polarization is regulated by extracellular matrix remodeling"

### Supplemental Figure 1

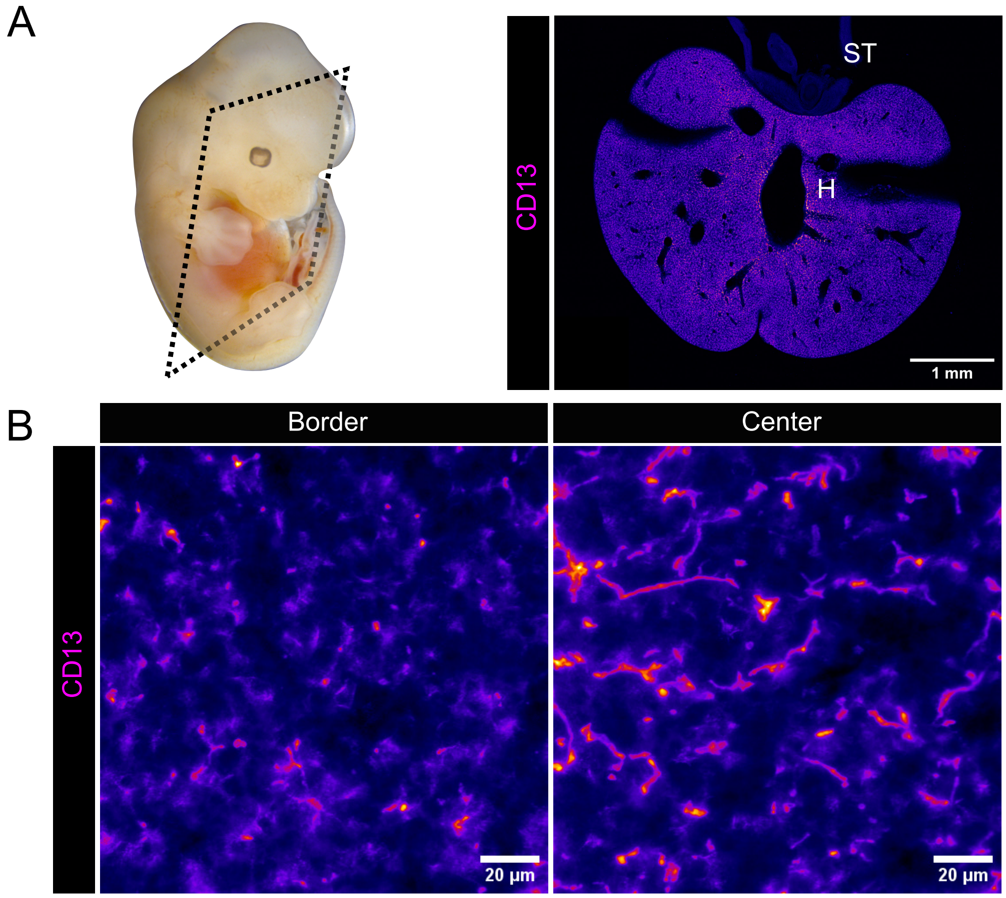


**Supplementary Figure1: Differences across the embryonic liver tissue.**

### Supplemental Figure 2

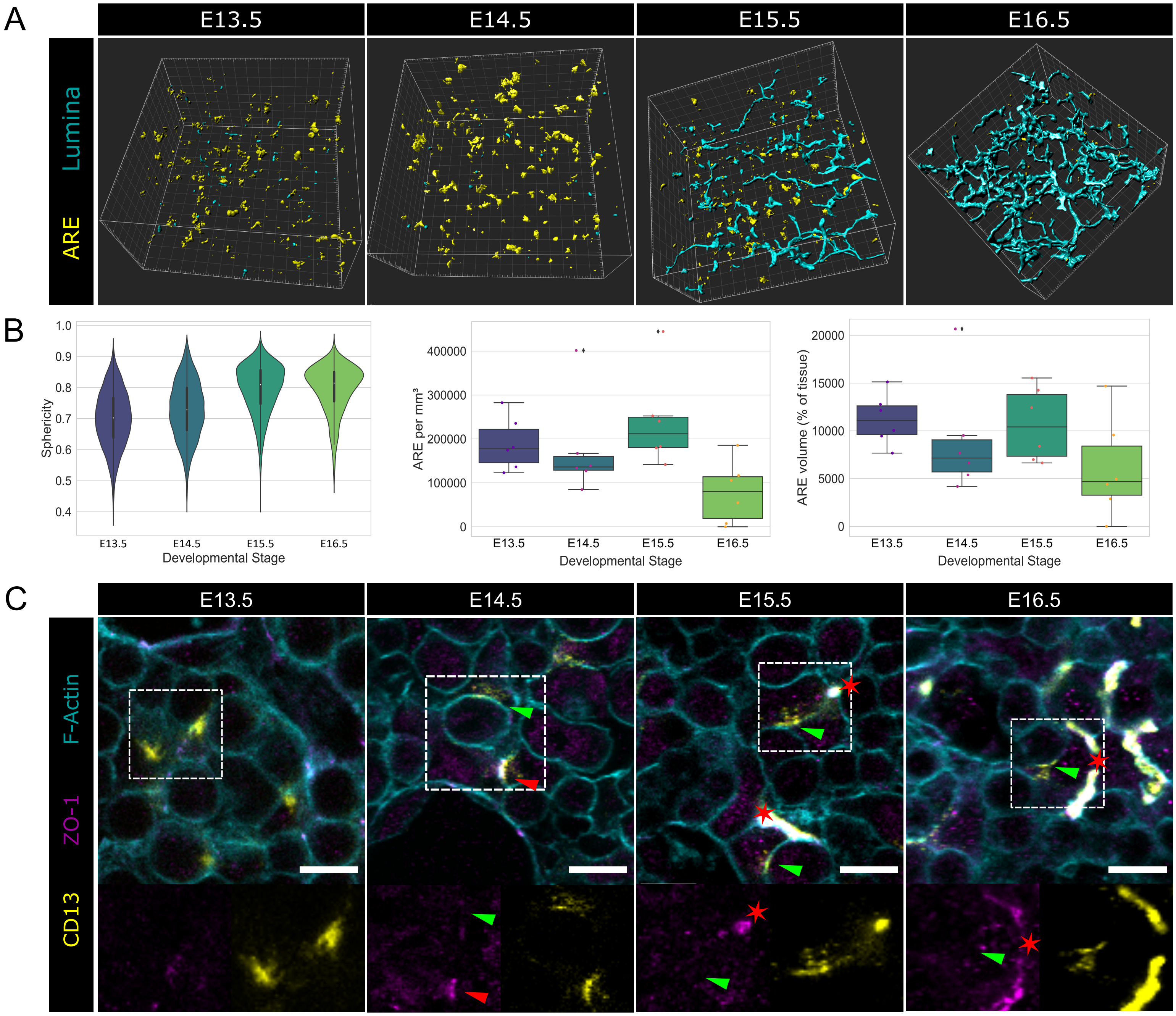


**Supplementary Figure2: ARE and lumen morphology.**

### Supplemental Figure 3

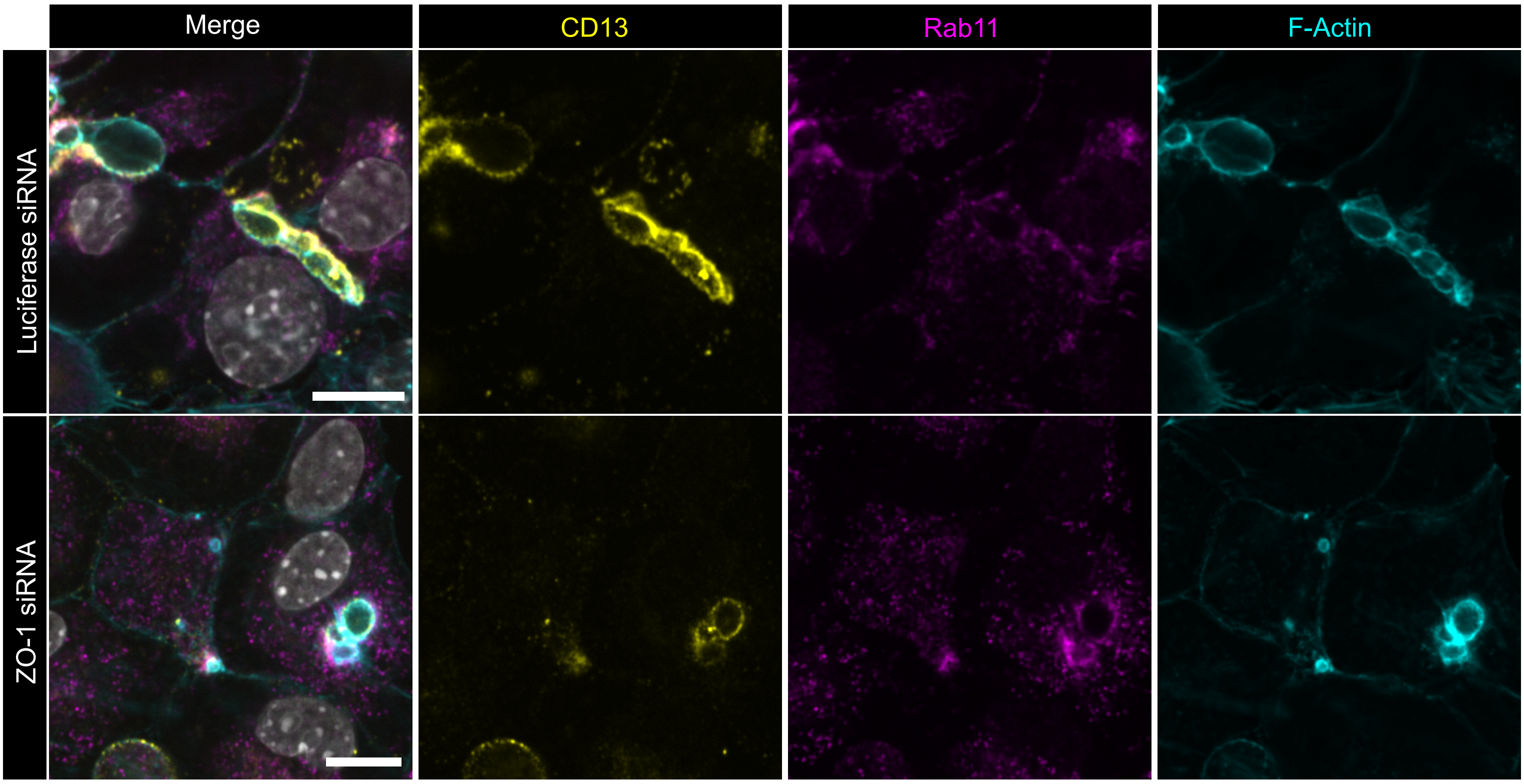

### Supplemental Figure 4

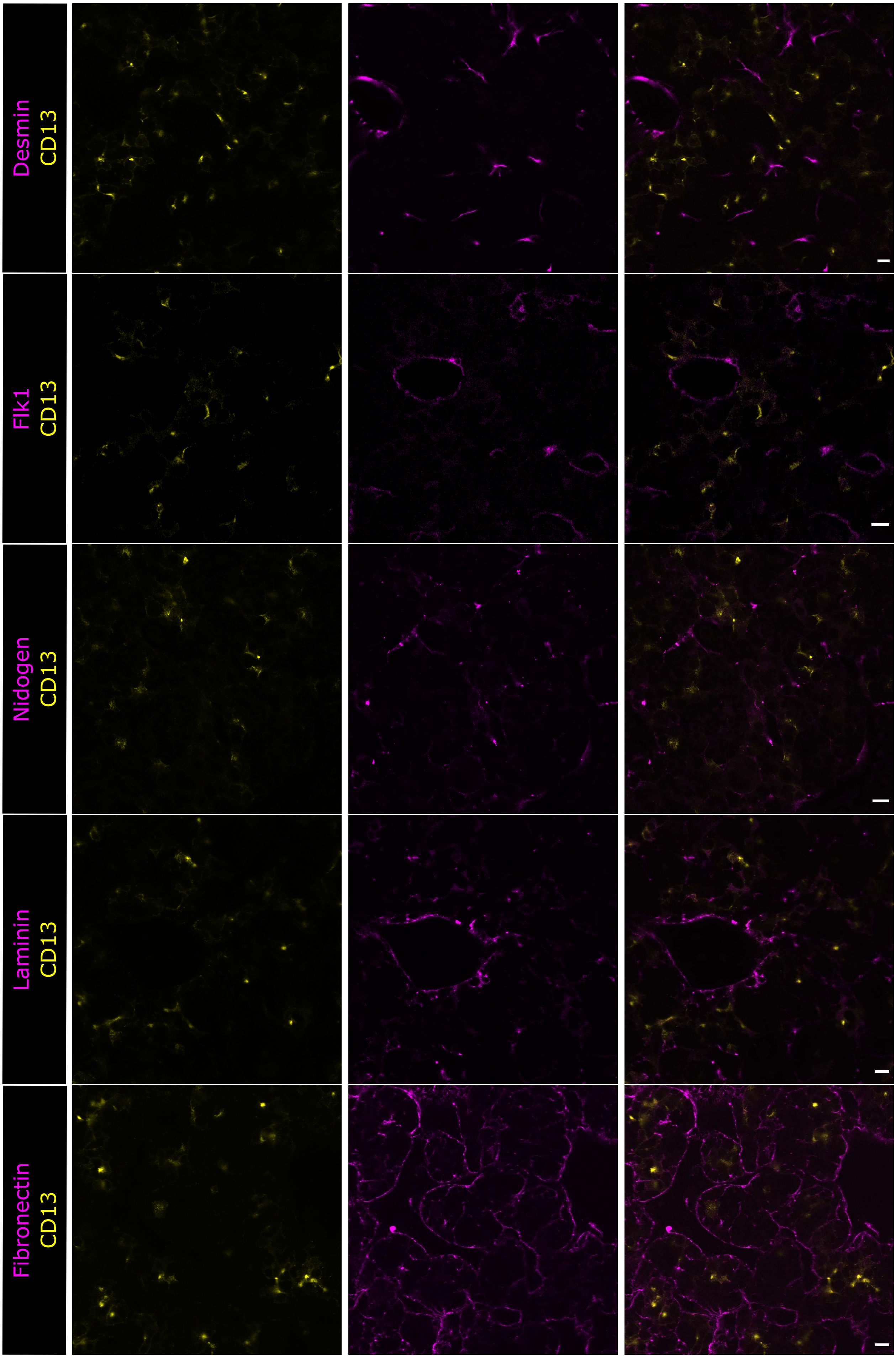


**Supplementary Figure4: Basal markers distribution**
