## Supplemental Figure 5 for "Hepatoblast iterative apicobasal polarization is regulated by extracellular matrix remodeling"

| Metalloprotease | GM6001 | CP 471474 | Marimastat | SD 2590 |
|---|---|---|---|---|
| 2 | | | | |
| 3 | | | | |
| 8 | | | | |
| 9 | | | | |
| 13 | | | | |
| 14 | | | | |
| 15 | | | | |
| Lumen formation | Yes | No | Yes | No |

**Supplementary Figure5: MMP inhibitor screen**
